# Alterations in background ECoG activity and behavioral deficits in a mouse model of CHD2-related developmental delay

**DOI:** 10.1101/2025.03.18.643778

**Authors:** Anat Mavashov, Shaked Turk, Yael Sarusi, Marina Brusel, Rotem Ben Tov Perry, Shir Quinn, Yael Almog, Karni Vilian, Igor Ulitsky, Moran Rubinstein

## Abstract

Heterozygous loss of function mutations in the *CHD2* gene, encoding for chromodomain helicase DNA-binding protein 2, are associated with severe childhood-onset epilepsy, global developmental delay, and autistic features. Here, we characterized the behavioral and epileptic phenotypes of a mouse model harboring a frameshift truncating mutation in the *Chd2* gene (*Chd2*^WT*/*m^ and *Chd2*^m/m^ mice). Genetic background dramatically affected the phenotypes. While no phenotypes were observed on the pure C57BL/6J background, crossing these mice onto the 129X1/SvJ genetic background gradually uncovered neurodevelopmental phenotypes. Transcriptomic analysis identified *Kcnj11* as a potential genetic modifier. On the 129X1/SvJ background, *Chd2*^m/m^ mice demonstrated growth retardation, and both *Chd2*^WT*/*m^ and *Chd2*^m/m^ showed motor deficits, including clasping behavior and reduced abilities to balance on a rotating rod. Autistic-like features were also observed, with *Chd2*^m/m^ showing reduced nest-building abilities and *Chd2*^WT*/*m^ demonstrating increased repetitive-like behavior in the marble burying test and altered social behavior. Quantitative analysis of electrocorticographic (ECoG) recordings revealed neuronal changes consisting of a global reduction in the total power of background activity in *Chd2*^WT*/*m^ and *Chd2*^m/m^ mice, as well as increased susceptibility to seizures induced by acute administration of 4-aminopyridine. Overall, this mouse model recapitulates multiple key phenotypes observed in CHD2 patients, providing a valuable platform to study the molecular basis and treatment options for this intractable disease.

## Introduction

Developmental disorders (DD) are a heterogeneous group of severe disorders with a strong genetic etiology. Over 40% of cases are caused by *de novo* mutations in hundreds of genes involved in synaptic and neuronal functions, transcriptional regulation, and chromatin remodeling (Satterstrom et al., 2020). Chromatin remodelers control chromatin structure and accessibility, regulating DNA replication, repair, and gene transcription. Among these, the Chromodomain Helicase DNA-binding (CHD) protein family, with its nine genes (*CHD1*– *CHD9*), is key in modulating gene expression essential for cellular differentiation and development. By their pivotal role, dysfunction of the CHD protein family has been implicated in cancers, and mutations in *CHD1*, *CHD2*, *CHD4,* and *CHD8* are associated with developmental disorders (Lamar and Carvill, 2018; Malone and Roberts, 2024).

*De novo* mutations in *CHD2* are identified in patients with developmental delay, mild to profound intellectual disability, often with seizures, photosensitivity, and autism spectrum disorder. Many of the pathogenic *CHD2* variants are truncating mutations due to frameshift, splicing, or deletions, indicating that C*HD2* haploinsufficiency is the cause of the disease (Galizia et al., 2015; Lamar and Carvill, 2018; Feng et al., 2022; Prince et al., 2024).

Efforts to develop animal models of *Chd2*-DD have included an early mouse model in which a gene trap vector was inserted in the 27^th^ intron of the *Chd2* gene, resulting in a genetic change expected to yield a protein that lacked the DNA binding domain (Marfella et al., 2006). These mice exhibited embryonic lethality when homozygous for the mutant allele, and growth defects as well as kidney abnormalities when heterozygous, but specific neuronal phenotypes were not described. Additionally, a zebrafish model with partial *Chd2* loss of function was generated, revealing a motor phenotype and seizure-like activity, providing insights into *CHD2* functions and associated phenotypes (Suls et al., 2013; Galizia et al., 2015). A more recent mouse model, with targeted replacement of the third *Chd2* exon by a stop cassette, demonstrated fewer GABAergic neurons and alteration in neuronal and synaptic functions without recapitulating clinical phenotypes (Kim et al., 2018) and mild, nonneuronal phenotypes on a homozygous background (Groza et al., 2023). Thus, while existing models of *Chd2*-DD have contributed to our understanding, none fully capture the phenotypes observed in patients.

Here, we report the characterization of a recently generated *Chd2* mouse model harboring a frameshift truncating mutation in the third exon (Rom et al., 2019). Examining these heterozygous and homozygous mutant mice demonstrates reduced *CHD2* expression, alteration in background electrocorticography (ECoG) oscillations, behavioral deficits, and an increased susceptibility to seizures. This model, which recapitulates several specific disease-related phenotypes, provides a valuable framework for understanding the *CHD2*-related mechanisms and defining specific readouts to explore the therapeutic effect of current and novel treatment options.

## Methods

### Animals

All animal experiments were approved by the Animal Care and Use Committee (IACUC) of Tel Aviv University. Mice used in this study were housed in a standard animal facility at the Goldschleger Eye Institute at a constant (22°C) temperature, on a 12-h light/dark cycle, with *ad libitum* access to food and water.

The *Chd2*^WT/m^ mice on the C57BL/6J (B6) background were generated at Weizmann Institute as described before (Rom et al., 2019). Briefly, the deletion of nucleotides 266 and 267 in the *Chd2* gene was induced by CRISPR/Cas9 with a single guide RNA. This deletion caused a frameshift mutation that, on the protein level, resulted in a change of four amino acids (starting from amino acid 90), converting the WT sequence KERI to GTDS*, with a nonsense mutation after amino acid 93 (Fig. 1A and Supplementary Fig. 1A).

**Fig. 1.**
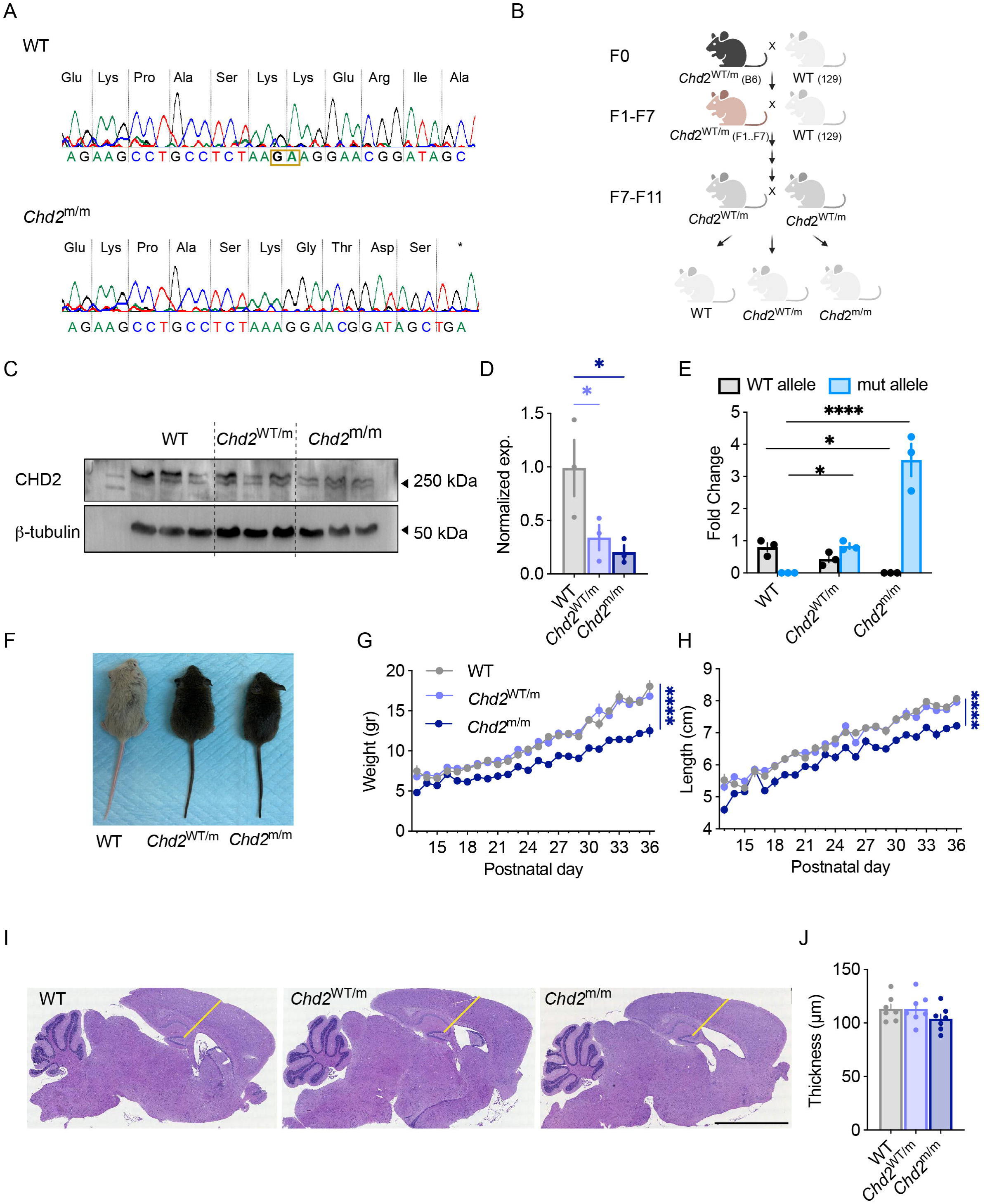
Reduced CHD2 expression and growth delay in *Chd2*^m/m^ mice. (A) The DNA and protein sequence of CHD2 in a WT mouse and *Chd2*^m/m^. The two nucleotides that were deleted are marked by a yellow box. (B) Depiction of the breeding scheme. *Chd2*^WT/m^ mice were crossed with WT mice on the pure 129x1/SvJ for at least seven generations prior to the breeding of the two *Chd2*^WT/m^ mice together to generate WT, *Chd2*^WT/m^ and *Chd2*^m/m^ mice. (C) Western blot analysis of CHD2 from whole brain extracts of three different mice from each genotype. Kilodaltons = kDa. (D) Quantification of CHD2 protein level after normalization to β-tubulin (n=3 for each genotype. Statistical analysis: One-way ANOVA followed by Holm-Sidak’s test). (E) Allele-specific quantification of *Chd2* transcripts (n=3 for each genotype. Statistical analysis: Two-way ANOVA followed by Holm-Sidak’s test). (F) Representative image of WT, *Chd2*^WT/m^, and *Chd2*^m/m^ at P35. (G-H) Reduced weight (G) and length (H) in *Chd2*^m/m^ mice (WT: n=40, *Chd2*^WT/m^: n=64, *Chd2*^m/m^: n=34. Statistical analysis: Mixed-model ANOVA followed by Tukey’s test). The normalized weight gain and the growth of males and females are depicted in Supplementary Fig. 4. (I-J) H&E-stained sagittal brain sections of WT, *Chd2*^WT/m^, and *Chd2*^m/m^ at 4 weeks old. The thickness of the hippocampal and cortical layers was measured using the yellow lines. Calibrator, 100 mm (WT: n=7, *Chd2*^WT/m^: n=7, *Chd2*^m/m^: n=7. Statistical analysis: One-way ANOVA followed by Holm-Sidak’s test).

*Chd2*^WT/m^ B6 mice (F0) were crossbred with WT 129X1/SvJ (129) mice (The Jackson Laboratory, Bar Harbor, ME, USA. Strain #000691) for one generation (Supplementary Fig. 2), four to six generations (Supplementary Fig. 3) or over seven generations (Figs. 1-3, 5). The transcriptomic analyses (Fig. 4) were done on mice from generations five and six. While crossing *Chd2*^WT/m^ mice on the 129 background, we observed difficulties in parental care, as many litters did not survive in the home cage with their biological parents. To address this, we paired a pregnant *Chd2*^WT/m^ mother with an experienced mother on a C57BL/6J background at a similar pregnancy stage. Together, the two mothers successfully raised both litters, and the pups were separated based on coat color (*Chd2* pups had white to brown coats and the B6 background had black coats) before further genotyping. PCR was performed using the primers and protocol described before (Rom et al., 2019).

All experiments were conducted on males and females, and the data were pooled together.

### Quantitative reverse-transcription PCR (qRT-PCR)

Reverse transcription was done using qScript Flex cDNA synthesis kit (Quanta Biosciences), using random primers. Quantitative PCR was performed in a ViiA 7 Real-Time PCR System (Thermo) in a 10 μl reaction mixture containing 0.1 mM forward and reverse primers, fast SYBR master mix (Applied Biosystems), and template cDNA. A reaction containing DDW instead of cDNA was used as a no-template control and was amplified for each primer pair. Actin or Gapdh was used as the reference gene for normalization.

The allele-specific primer pairs used for mRNA expression level analysis were:

F_Chd2-WT W2641 TGAGAGTGAATCGGCAGGTT

R_Chd2-WT W2642 CACATCAGCTATCCGTTCCTTC

R_Chd2-mut W2643 CACATCAGCTATCCGTTCCTTT

F_Kcnj11 TGGGCACGTGGAAAGTGAAG

R_Kcnj11 CACAGGGGTAAACCAGTCCC

F_Actin TTGGGTATGGAATCCTGTGG

R_Actin CTTCTGCATCCTGTCAGCAA

F_Gapdh GTCGGTGTGAACGGATTTG

R_Gapdh GAATTTGCCGTGAGTGGAGT

### Western blot

Total protein was extracted from whole brains by lysis with RIPA supplemented with protease inhibitors and DTT 1 mM. Proteins were resolved on 8% SDS-PAGE gels and transferred to a polyvinylidene difluoride (PVDF) membrane. After blocking with 5% nonfat milk in PBS with 0.1% Tween-20 (PBST), the membranes were incubated with the primary antibody, followed by the secondary antibody conjugated with horseradish peroxidase. Primary antibodies were as follows: anti-Chd2 (Novus, NBP2-92115-0.1ml, 1:1,000 dilution), anti-β-tubulin (Sigma, #T4026, 1:1,000 dilution). Secondary antibodies were anti-rabbit (#111-035-003, 1:10,000 dilution) and anti-mouse (#115-035-003, 1:10,000 dilution). Blots were imaged on an Azure Imager system and were quantified with ImageJ software.

### H&E staining

Brains were fixed in 4% paraformaldehyde in 1x PBS (4% PFA/PBS) overnight at 4°C. Following fixation, tissue samples were washed in 1x PBS (3 x 20 minutes each) to remove residual 4% PFA/PBS and dehydrated through the series of graded ethanol, isopropanol, and toluene to displace the water. Processed tissues were then embedded in paraffin in the plastic cassettes and allowed to set. Paraffin blocks were stored at room temperature until further use. Serial sagittal sections of 10 μm in thickness were cut on microtome through the entire brain. The sections were first de-waxed using X-TRE-SOLV washes (3 x 5 minutes each), subsequently re-hydrated in a decreasing series of ethanol (100%, 95% 70%, 1 x 5 minutes each), sections were then washed in distilled water (1 x 2 minutes each). The staining protocol was as follows: Haematoxylin solution (Leica, cat #3801521E) for 5 minutes, rinsed quickly in distilled water, wash in 70% ethanol (1 x 3 minutes) and then stain with Eosin (Sigma, cat #HT110332-1L) for 1.2 minutes, differentiated in 95% ethyl alcohol and dehydrate quickly in 100% alcohol, then cleared using X-TRE-SOLV washes (2 x 3 minute each) and mounted. The sections were imaged using Leica DM4000 B microscope with Leica DFC365 FX CCD camera and Leica application suite (LAS) X software. The thickness of hippocampal and cortical layers was quantified using Fiji ImageJ.

### Behavioral tests

Behavioral experiments were conducted on three genotypes: WT, *Chd2*^WT/m^ and *Chd2*^m/m^. Mice were handled for 5 minutes before testing and habituated to the experimental room for 30 minutes on the test day. The experimental apparatus and objects were cleaned with 70% ethanol, rinsed with water, and dried between trials.

### Hindlimb clasping

The mice were held upside down by the base of their tail and videotaped for 30 seconds (LifeCam HD-6000, Microsoft, Redmond, WA, USA). Clasping severity was scored from 0 to 3: 0=hindlimbs splayed outward and away from the abdomen (normal); 1=one hindlimb retracted inward toward the abdomen for at least 50% of the observation period; 2=both hindlimbs partially retracted inward toward the abdomen for at least 50% of the observation period; 3=both hindlimbs completely retracted inward toward the abdomen for at least 50% of the observation period. Scores of 0.5 were utilized when appropriate (Guyenet et al., 2010; Sampson et al., 2016).

### Open field

Open field testing was performed as previously described (Fadila et al., 2020). Briefly, the mice were placed in the center of a square Plexiglas apparatus (50 cm x 50 cm) for 10 minutes. To assess anxiety-like behavior, two zones were defined within the arena: the “outer” zone, which included all areas within 10 cm of the walls, and the “inner” zone, comprising the remaining central area. Live tracking was achieved via a monochrome camera (Basler acA1300-60gm, Basler AG, Ahren, Germany) connected with EthoVision XT 13 software (Noldus Technology, Wageningen, Netherlands). Some mice, probably due to the poor motivation for spontaneous exploratory behavior of 129X1/SvJ mice (Tang and Sanford, 2005), were reluctant to explore the arena. Therefore, mice that traveled less than 500 cm and did not enter the inner zone were excluded from the analysis.

### Rotarod

Two protocols were tested: *i*) The rotarod test was conducted over three consecutive days, with five trials per day, each separated by at least 30 minutes (Fig. 2D). On the first day, mice were habituated to the rod by being placed on a stationary rod for 5 minutes before the first trial. During each trial, the rod accelerated from 4 to 40 RPM over 300 seconds. The performance score for each trial was either the duration the mouse remained on the rod or 300 seconds, depending on which occurred first. *ii*) Mice were placed on a rotating rod accelerating from 4 to 40 RPM over the course of 180 seconds, and the time at which each mouse fell was recorded. Each mouse had five consecutive trials. The final score was calculated as the average of the 3 longest trials (Supplementary Fig. 5A).

**Fig. 2.**
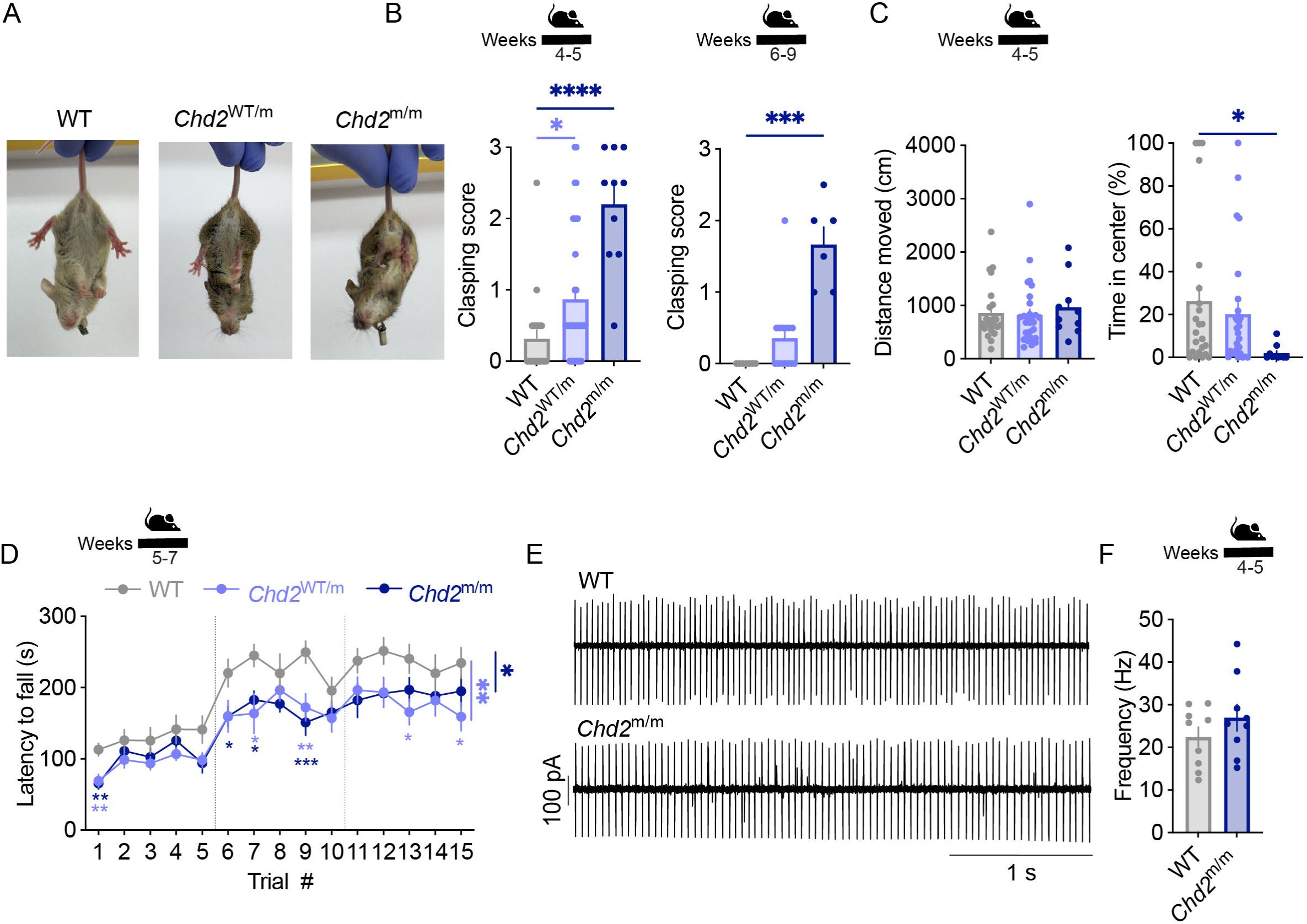
Motor deficits in *Chd2* mutant mice. (A) *Chd2* mutant mice show clasping when suspended from their tail. (B) The average clasping score of juvenile mice (left, WT: n=22, *Chd2*^WT/m^: n=34, *Chd2*^m/m^: n=10. Statistical analysis: Kruskal-Wallis test followed by Dunn’s test) and adult mice (right, WT: n=7, *Chd2*^WT/m^: n=14, *Chd2*^m/m^: n=6. Statistical analysis: Kruskal-Wallis test followed by Dunn’s test). (C) The total distance traveled in the open field (left) and the percent of time spent in the center of the arena (right) (Juvenile mice. WT: n=23, *Chd2*^WT/m^: n=28, *Chd2*^m/m^: n=10. Statistical analysis: Kruskal-Wallis test followed by Dunn’s test). The results of adult mice are depicted in Supplementary Fig. 5C. (D) The ability to balance on the rotarod was tested 5 times in 30-minute intervals for 3 consecutive days (WT: n=12, *Chd2*^WT/m^: n=13, *Chd2*^m/m^: n=12. Statistical analysis: Two-way ANOVA followed by Dunnett’s test). (E) Example traces of spontaneous firing of PNs. (F) Average firing rates (WT: n=9 cells from 3 mice, *Chd2*^m/m^: n=9 cells from 3 mice. Statistical analysis: Unpaired t-test).

### Four limb-hanging test

This test assesses motor deficits in mice (Kim et al., 2010). The procedure consisted of two 60-second trials separated by a 30-minute interval. In each trial, the mouse was placed upside down, holding a grid, and the time it remained on the grid, up to a maximum of 60 seconds, was recorded. The final score was calculated as the average of the two trial durations (Supplementary Fig. 5B).

### Nest building test

Approximately one hour before the dark phase, mice were placed individually in home cages with clean bedding, 1.2 g of pre-weighed dense cotton, and no enrichment. After ∼16 hours, nests were evaluated based on two metrics: the percentage of untorn cotton (brushed-off bedding) relative to the initial weight and the nest score on Deacon’s 5-point scale (Deacon, 2006). Scores were assigned as follows: 1=cotton not noticeably touched (>90% intact); 2=cotton partially torn (50–90% intact); 3=cotton mostly shredded (>50%) but with no identifiable nest site; 4=cotton mostly shredded (>90%), with an identifiable, but flat nest; 5=cotton mostly shredded (>90%), with an identifiable, heightened nest. Half-scores were assigned for performances between severity levels.

### Marble burying

The mice were placed in a cage (36 cm L x 26 cm W x 18 cm H) with a 5 cm layer of SaniChip bedding, and twenty glass marbles (1.4 cm in diameter, varying in color) which were arranged in an evenly spaced grid formation (4 rows x 5 columns). Mice were placed in a corner of the cage and videotaped for 30 minutes using a webcam. Two parameters were assessed: the number of buried marbles (defined as those covered by bedding to at least 2⁄3 of their height) and the total time spent digging. Video recordings were analyzed manually offline using EthoVision XT 13 tracking software (Noldus Technology, Wageningen, Netherlands).

### Social interaction

Two tests were conducted: *i*) Interaction between pairs of mice of the same genotype (Fig. 3F). The test was conducted in a square open-field arena (50 × 50 cm), allowing the mice to rest at the corners as needed. Each pair consisted of stranger mice with the same genotype, in addition to matching sex, age, and genetic background, ensuring that each pair represented a single statistical unit. One day before the experiment, each mouse was placed alone in the arena for 15 min to acclimate the arena. On the following day, pairs of mice were placed together in the arena for another 15 min. Their activity was recorded (Basler acA1300-60gm, Basler AG, Ahren, Germany) and analyzed offline using EthoVision XT 17 with the Social Interaction add-on module (Noldus Technology, Wageningen, Netherlands). The trials were divided into 5-minute time bins for analysis; for each bin, we calculated the percent of interaction time (defined as proximity of 9.5 cm or below) and the mean distance between the mice. *ii*) Interaction with a WT mouse (Supplementary Fig. 5D, E). The test was conducted in a circular open-field arena (50 cm diameter) with an experimental mouse and an unfamiliar WT interacting mouse of the same sex, color, age, and size. Each mouse was acclimated to the arena individually for 3 minutes before the trial. The pairs of mice were placed together in the arena for 20 minutes, and their behavior was recorded. The recordings were analyzed manually using EthoVision XT 13 (Noldus Technology, Wageningen, Netherlands).

**Fig. 3.**
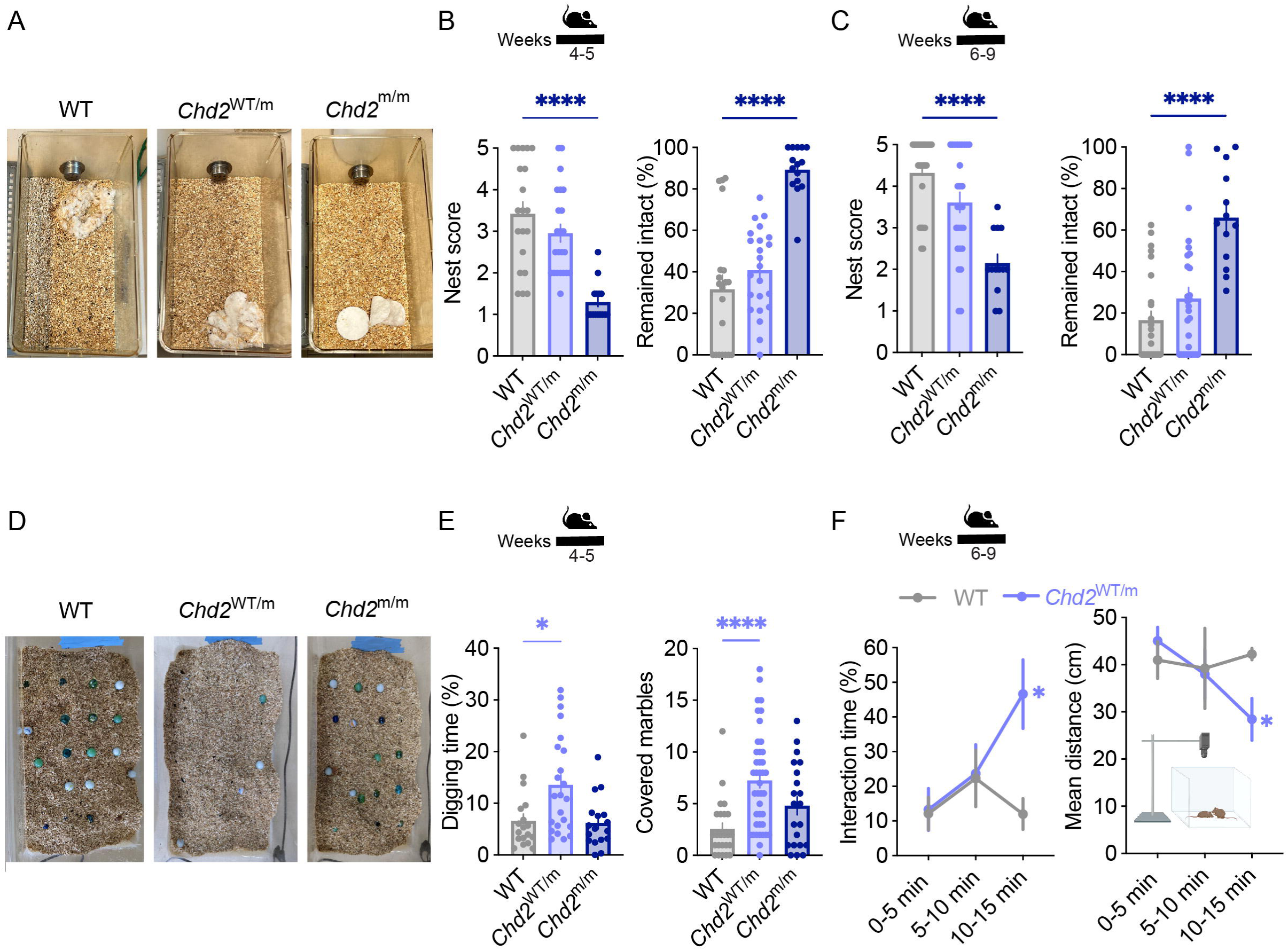
Autistic-like features in *Chd2* mutant mice. (A) Representative images of nests built by juvenile mice. (B, C) Nest complexity scores and the percent of remaining nesting materials (Juvenile mice (B), WT: n=20, *Chd2*^WT/m^: n=23, *Chd2*^m/m^: n=15. Adult mice (C), WT: n=25, *Chd2*^WT/m^: n=27, *Chd2*^m/m^: n=13. Statistical analysis: Kruskal-Wallis test followed by Dunn’s test). (D) Representative images of covered marbles. (E) The percent digging time and the number of covered marbles (Juvenile mice. WT: n=24, *Chd2*^WT/m^: n=41, *Chd2*^m/m^: n=21. Statistical analysis: Kruskal-Wallis test followed by Dunn’s test). (F) Cumulative social interaction (left) and the mean distance between pairs of stranger mice from the same genotype in 5 minute bins (WT: n=10 mice in 5 pairs, *Chd2*^WT/m^: n=24 mice in 12 pairs. Statistical analysis: Two-way ANOVA followed by Holm-Sidak’s test).

### Cerebellum brain slice recordings

Cerebellum brain slices were prepared as described before (Fadila et al., 2020). Briefly, 300 μm sagittal cerebellum slices were cut using a Leica VT1200S vibratome (Leica Biosystems, Wetzlar, Germany). Loose-patch (15-100 MΩ seal resistance) extracellular recordings were made from the soma of visually identified Purkinje neurons (SOM; Sutter Instrument, Novato, CA, USA). Only one cell was recorded from each lobe, and no more than two cells from each slice. The electrode pipettes (resistance of 2 - 4 MΩ) were filled with ACSF, which was also the extracellular solution (125 mM NaCl, 3 mM KCl, 2.0 mM MgCl_2_, 2.0 mM CaCl_2_, 1.25 mM NaH_2_PO_4_, 26 mM NaHCO_3_, and 10 mM D-glucose, saturated with 95% O_2_ and 5% CO_2_). Recordings were obtained at room temperature using a Multiclamp 700B amplifier (Molecular Devices, San Jose, CA, USA) and Clampex 10.7 software (Molecular Devices) under a voltage clamp at zero current.

### RNA extraction and sequencing

Total RNA was extracted from the hippocampus, cortex, cerebellum, and thalamus using the TRI REAGENT (MRC) according to the manufacturer’s protocol. Strand-specific mRNA-seq libraries were prepared from 1 µg total RNA using the CORALL-mRNA-Seq-V2 (Lexogen), according to the manufacturer’s protocol, and sequenced on a Novaseq 6000 machine to obtain 75 nt and 150 nt single or paired-end reads.

### RNA-seq data analysis

After adaptor trimming, RNA-seq reads were mapped to the mm10 mouse genome assembly using STAR (Dobin et al., 2013) and gene expression was quantified using mouse RefSeq gene annotations and RSEM (Li and Dewey, 2011), and differential expression between genotypes was computed using DeSeq2 (Love et al., 2014).

### RNAscope FISH

Brains were flash frozen on dry ice in a tissue-freezing medium and sliced on a cryostat (Leica CM 1950) into 10-mm sections, adhered to SuperFrost Plus slides (VWR), and immediately stored at -80°C until use. Samples were processed according to the ACD RNAscope Fluorescent Multiplex Assay manual using *Kcnj11* probes cat No. 704611, and Abcc8 probes cat No. 498131. Imaging was performed on a Nikon-Ti-E inverted fluorescence microscope with a 1003 oil-immersion objective and a Photometrics Pixis 1024 CCD camera using MetaMorph software. RNAscope analysis was done using IMARIS (v7.7.2) software.

### Electrocorticography (ECoG) recordings and drugs administration

Electrode implantation and video-ECoG recordings were performed as previously described (Fadila et al., 2020; Mavashov et al., 2023). Recordings lasted 2-4 hours between 8:00 AM and 6:00 PM. The first 30 minutes of the recording were considered acclimation periods and excluded from the analysis. ECoG signal pre-processing and further analysis were conducted as described (Quinn et al., 2023).

To assess potential therapeutic effects, Glibenclamide (GBC, 1mg/kg in saline with 30% Polyethylene glycol 400 and 0.1% DMSO; Alomone Labs, Jerusalem, Israel) was administered as a single intraperitoneal injection after 2 hours of baseline ECoG recording. Following the injection, recording continued for an additional 2 hours. The dosage was selected to achieve therapeutic-relevant concentration following acute administration.

To induce seizures, 4-aminopyridine (4-AP, 4 mg/kg in saline; Sigma-Aldrich, Saint Louis, MO, USA) was injected intraperitoneally after at least 1 hour of baseline ECoG recording. Elevation of the body core temperature started 30 min after 4-AP administration in mice that did not show epileptic activity and was done as previously described (Almog et al., 2021). Briefly, the body temperature was increased by 0.5°C every 2 min using a heat lamp attached to a feedback temperature controller (TCAT-2DF, Physitemp Instruments) until presentation of epileptic activity. The temperature was not increased above 42 °C.

### Statistical analyses

Data are reported as mean ± standard error (SE). All statistical tests were carried out using GraphPad Prism (Boston, MA, USA), and details of the specific test are listed in the Figure legend. Differences were considered significant at p < 0.05 and marked on the Figure: p < 0.05, *; p < 0.01, **; p < 0.001, ***; p < 0.0001, ****. The specific p values, as well as the number of mice, males and females, are listed in Supplemental Table 1.

## Results

### Growth retardation, hindlimb clasping and hyperactivity in *Chd2* mice on enriched 129X1/SvJ genetic background

*De novo* frameshift mutations in CHD2-DD were identified in ∼25% of the CHD2 patients (De Maria et al., 2022). This genetic change was recapitulated in a mouse using CRISPR-mediated deletion of nucleotides 266 and 267 in the *Chd2*, resulting in the alteration of four amino acids in the N-terminal domain of the protein. This change converted the wild-type (WT) amino acid (a.a) sequence of KERI to GTDS* leading to reduced CHD2 expression (Fig. 1A and Supplementary Fig. 1). This original model was created in the C57BL/6J background, presented no obvious neurological phenotypes, growth delay, or behavioral seizures, and had a normal lifespan (Rom et al., 2019).

The most characteristic phenotype in CHD2-DD is epilepsy and seizures (Lamar and Carvill, 2018; Prince et al., 2024). In animal models, the detection of epileptic activity provides an unequivocal objective demonstration of a disease-relevant phenotype and a potential robust biomarker to examine disease progression and the therapeutic effect of treatments (Wang and Frankel, 2021; Barker-Haliski and Hawkins, 2024). To examine the presentation of epileptic activity in *Chd2* mutant mice on the C57BL/6J background, ECoG recordings of heterozygous (*Chd2*^WT/m^) and homozygous (*Chd2*^m/m^) mutant mice were performed. However, in line with previous reports (Kim et al., 2018), no epileptic activity was observed (Supplementary Fig. 2A).

Due to this, we wondered if a different genetic background would reveal CHD2 disease-relevant phenotypes, given that the genetic strain background of mice was shown to affect the phenotypic presentation of neurological disorders (McGraw et al., 2017). To that end, *Chd2* mutant mice were crossed with 129X1/SvJ mice for one generation, generating *Chd2*^WT/m^ on a 50:50 C57BL/6J:129X1/SvJ background, followed by a crossbreeding of *Chd2*^WT/m^ pairs to generate WT, *Chd2*^WT/m^, and *Chd2*^m/m^ mice. Examination of the locomotor exploratory behavior of these mice in a novel arena using the open field test did not indicate increased anxiety or hyperactivity (Supplementary Fig. 2B, C), which is often detected in mouse models of DD (Silverman et al., 2010).

Moreover, the vast majority of the recorded *Chd2* mutant mice (83%) had no signs of epilepsy or changes in the properties of ECoG activity (Supplementary Fig. 2D-F). Despite that, in one of the recorded *Chd2*^m/m^ mice (1 of 6), we did notice spontaneous epileptic activity (Supplementary Fig. 2G). This single observation prompted us to continue the crossing onto the 129X1/SvJ background.

Additional studies were performed after four to six generations of crossing onto the 129X1/SvJ background (>90% 129X1/SvJ, Supplementary Fig. 3A). Two age groups were studied: juvenile mice (4-5 weeks old) and adult mice (6-8 weeks old). Mice in the 4-5 week range are weaned from their parents but have not yet reached sexual maturity (Dutta and Sengupta, 2016). In humans, this developmental stage is most related to toddlers and preschoolers, which coincides with the typical onset of CHD2-related epilepsy (Thomas et al., 2015; De Maria et al., 2022; Zhu et al., 2022). Sexual maturation is achieved in mice by 6 weeks of age, after which they are considered adults. This stage is roughly analogous to the teenage to young adult years in humans. Examination of the exploratory behavior in the open field of these *Chd2* mutant mice on enriched 129X1/SvJ background demonstrated increased locomotion in adult *Chd2*^m/m^ mice (Supplementary Fig. 3B). Furthermore, ECoG recordings revealed reduced power of the background activity in these mice (Supplementary Fig. 3C-J). The detection of these behavioral and electrographic phenotypes in adult *Chd2*^m/m^ mice suggests that the genetic background in mice modulates the presentation of the CHD2-related phenotype, and enrichment of the 129X1/SvJ background uncovers some of these phenotypes.

### Growth delay and motor deficits in *Chd2* mutant mice

Further examinations were conducted in mice crossed onto the 129X1/SvJ background for at least seven generations (>99% 129X1/SvJ background, Fig. 1B).

Analysis of the CHD2 protein levels in whole brain lysates demonstrated the expected ∼50% and ∼75% reduction in the CHD2 in *Chd2*^WT/m^ and *Chd2*^m/m^ mice, respectively, compared to WT littermates (Fig. 1C, D). Residual protein expression in *Chd2*^m/m^ is possibly related to an alternative start codon that the mutation puts back into frame with most of the protein (Supplementary Fig. 1A). Examination of the mRNA expression of the WT *Chd2* allele showed a trend for reduction in the expression of this allele in *Chd2*^WT/m^ and *Chd2*^m/m^ mice, together with increased expression of the mutant allele. The mutant allele showed a 3-fold increase in the *Chd2*^m/m^ mice, indicating that it was not subject to nonsense-mediated decay, and that some compensatory mechanism leads to an increase in *Chd2* mRNA expression when most CHD2 protein is lost (Fig. 1E).

Through their development, *Chd2*^m/m^ weighed less and were shorter in length, while the development of *Chd2*^WT/m^ and WT mice was comparable (Fig. 1F-H). Nevertheless, histological analyses did not demonstrate alterations in gross brain morphology or the thickness of the hippocampal and cortical layers (Fig. 1I, J).

Next, we tested the mice for DD-related behavioral phenotypes. Hindlimb clasping was observed in *Chd2*^WT/m^ and, more notably, in *Chd2*^m/m^ mice (Fig. 2A, B). The locomotor activity in the open field was similar between WT and *Chd2* mutant mice, but juvenile *Chd2*^m/m^ spent less time in the center of the arena, suggesting increased anxiety (Fig. 2C and Supplementary Fig. 5C). On the rotarod, *Chd2* mutant mice showed reduced abilities to balance when tested five times a day over the course of three consecutive days, indicating impaired motor abilities (Fig. 2D). Motor deficits are often associated with alteration in the activity of Purkinje neurons (PNs) in the cerebellum, and were observed in multiple forms of neurodevelopmental disorders, including epilepsy and autism spectrum disorders (Shevelkin et al., 2014; Cook et al., 2021). However, the firing rate of PNs from WT mice and *Chd2*^m/m^ mice were similar in extracellular recordings (Fig. 2E, F), indicating that the observed motor deficits are not associated with changes in spontaneous firing of PNs. Together, these data demonstrate growth retardation and impaired motor functions in *Chd2* mutant mice.

### Autistic-like features in *Chd2* mutant mice

Autistic traits are frequent in *CHD2* patients (Lamar and Carvill, 2018; Prince et al., 2024). Nesting is an innate rodent behavior that contributes to their well-being, and its impairment was reported in multiple rodent models of developmental and psychiatric disorders (Jirkof, 2014). In *Chd2*^m/m^ mice, we observed reduced nesting abilities, with less consumption of nesting materials and lower nest complexity (Fig. 3A-C), indicating an impairment in this daily function. In addition, in the marble burying test, which correlates with repeated behavior (Silverman et al., 2010), *Chd2*^WT/m^ mice spent more time digging and covered more marbles (Fig. 3D-E).

To further explore the social behavior of these mice, we examined the interaction time between an interacting WT mouse and a tested WT or mutant *Chd2* mouse when placed in a novel arena (Silverman et al., 2010). This test demonstrated a similar interaction time between the genotypes (Supplementary Fig. 5D, E). However, as the interacting WT mouse may have affected the behavior of the tested mouse, we also examined the interaction between two stranger mice of the same genotype. Using this paradigm, we uncovered altered social behavior in *Chd2*^WT/m^, consisting of increased interaction time and increased proximity between the mice during the last 5 minutes of the 15 minute test (Fig. 3F).

Together, these findings highlight behavioral deficits in *Chd2* mutant mice, consistent with the rodent manifestation of autistic-like features.

### Differential expression of *Kcnj11* in *Chd2* mutant mice

Following the uncovering of electrographic and behavioral changes in *Chd2* mutant mice on the 129x1/SvJ background, we sequenced RNA from different brain regions in an attempt to expose trancriptomic changes and identify genes that may have contributed to the exacerbation of disease-related phenotypes on the 129X1/SvJ background.

Focusing on ion channels and epilepsy-related genes that were differentially expressed in *Chd2* mutant mice, we identified an increased expression of *Kcnj11* mRNA in multiple brain regions in *Chd2*^WT/m^ and *Chd2*^m/m^ (Fig. 4A). The upregulation of *Kcnj11* was also confirmed using qPCR from whole brains. Interestingly, while *Kcnj11* was upregulated in *Chd2* mutant mice on the 129X1/SvJ, it was downregulated in *Chd2* mutant mice on the pure C57BL/6J background (Fig. 4B).

**Fig. 4.**
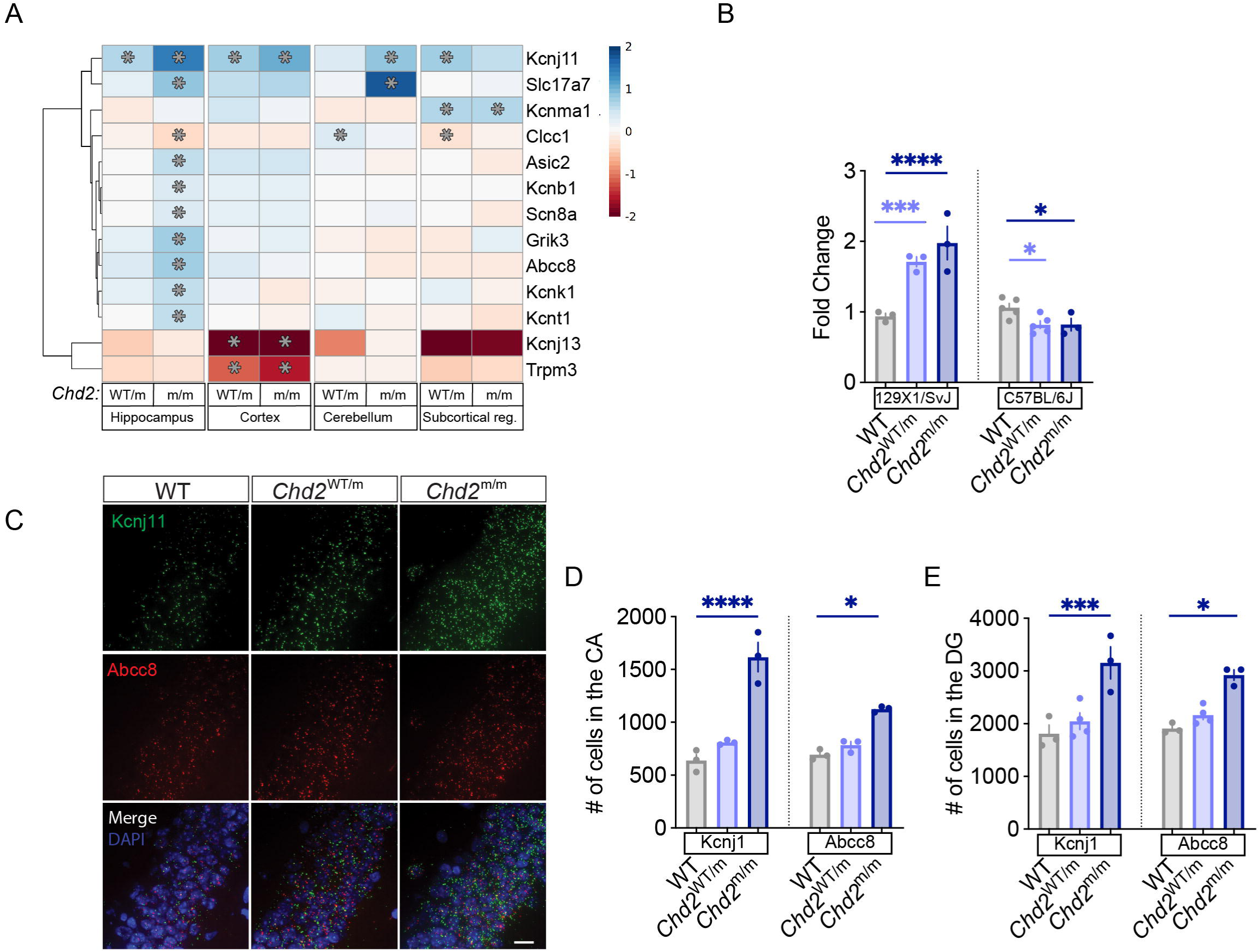
Upregulation of *Kcnj11* in *Chd2* mutant mice. (A) Heatmap showing relative expression of ion-channels-related genes that exhibited a significant change in gene expression in the hippocampus, cortex, cerebellum, and subcortical regions of 4-weeks-old *Chd2*^WT/m^ and *Chd2*^m/m^ mice compared to WT littermates (WT: n=3, *Chd2*^WT/m^: n=3, *Chd2*^m/m^: n=4. \**p*<0.05). (B) qPCR qunatification of *Kcnj11* from whole brains of WT and *Chd2* mutant on the 129X1SvJ and C57BL/6J backgrouds (129X1/SvJ, WT: n=3, *Chd2*^WT/m^: n=3, *Chd2*^m/m^: n=3. C56BL/6J, WT: n=5, *Chd2*^WT/m^: n=5, *Chd2*^m/m^: n=3. Statistical analysis: Two-way ANOVA followed by Holm-Sidak’s test.(C) RNAscope with probes targeting *Kcnj11* (green), *Abcc8* (red), and DAPI staining (blue) in the coronal section of the CA1 region of the hippocampus. Scale bar 20μm. (C-D) Quantification of *Kcnj11*^+^ and *Abcc8*^+^ cells throughout the CA (C) and the DG (D) regions in WT, *Chd2*^WT/m^ and *Chd2*^m/m^ (WT: n=3, *Chd2*^WT/m^: n=3 or 4, *Chd2*^m/m^: n=3. Statistical analysis: Kcnj11: One-way ANOVA followed by Holm-Sidak’s test; Abcc8: Kruskal-Wallis test followed by Dunn’s test).

In the membrane, KCNJ11 assembles with SUR1, encoded by the *Abcc8* gene, to form the ATP-sensitive potassium (K_ATP_) channel. The mRNA expression of the *Abcc8* was also increased in the hippocampus of *Chd2*^m/m^ mice (Fig. 4A). Next, we performed FISH using RNAscope method with probes targeting *Kcnj11* and *Abcc8* in the hippocampus of WT and *Chd2* mutant mice. In agreement with the RNA-seq data, the number of *Kcnj11* and *Abcc8* positive cells with the stratum pyramidal layer of cornu ammonis (CA) regions and the granular layer of the dentate gyrus (DG) was higher in *Chd2^m/m^* mice compared to WT mice (Fig. 4B-D). Thus, on the 129x1/SvJ background, there is a correlation between *Kcnj11* upregulation and the *Chd2* mutation, indicating that this may be a CHD2 target dysregulated in the brain.

### Altered background ECoG oscillations and 4-AP induced epileptic activity in *Chd2* mutant mice

The additional behavioral phenotypes revealed with further crossing onto the 129X1/SvJ background prompted us to repeat the ECoG recordings. Spontaneous epileptic activity was not detected in any of the recorded mice. However, we did observe a marked reduction in the power of background oscillations with lower power in *Chd2*^WT/m^ and *Chd2*^m/m^ mice, observed across multiple frequency bands (Fig. 5A-D). Further exploration of the relative power distribution within the ECoG signal demonstrated a lower contribution of the theta frequency band and an augmented contribution of the gamma frequency band in *Chd2*^m/m^ (Fig. 5E). Altered interhemispheric phase synchrony, or coherence, of background oscillations, was found in various brain disorders, including epilepsy and autism (Bowyer, 2016), as well as in *Chd2* mutant mice (Kim et al., 2018). Reduced coherence was also seen in *Chd2*^WT/m^ mice, mainly in the alpha and beta frequency bands (Fig. 5F).

**Fig. 5.**
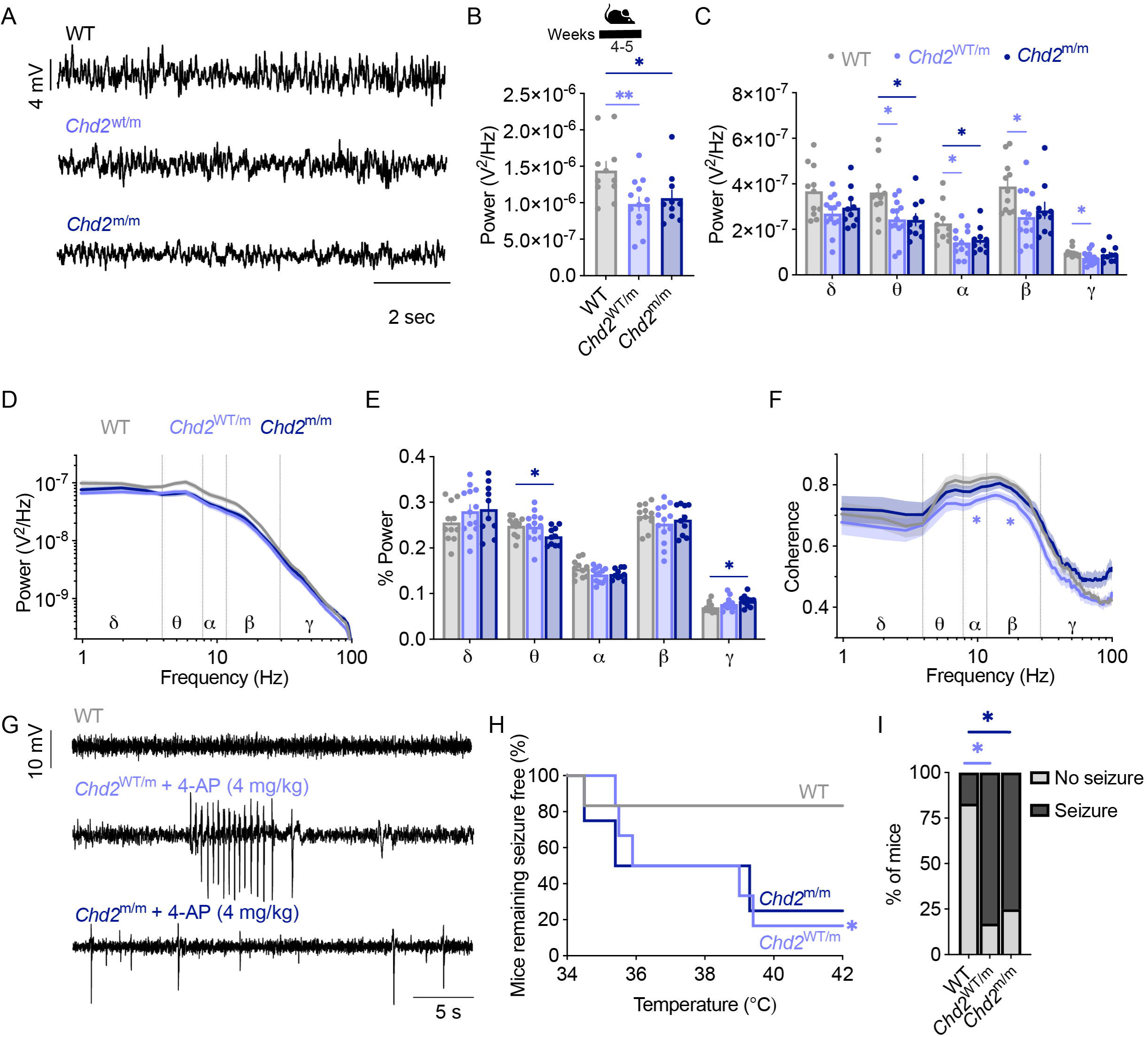
Reduced ECoG power and 4-AP induced epileptic activity in *Chd2* mutant mice. (A) Examples of background ECoG activity. (B) Total power (WT: n=11, *Chd2*^WT/m^: n=13, *Chd2*^m/m^: n=10. Statistical analysis: One-way ANOVA followed by Dunnett’s test). (C) Total power in the following frequency bands: δ, 0.9–3.9 Hz; θ, 4.8–7.8 Hz; α, 8.7–11.7 Hz; β, 12.6–29.2 Hz; γ, 30.2–99.6 Hz. Statistical analysis: Two-way ANOVA followed by the Holm-Sidak’s test. (D) Power spectral density. (E) The relative power in the different frequencies (The sum of the power in each frequency band divided by the total power). Statistical analysis: Two-way ANOVA followed by the Holm-Sidak’s test. (F) Interhemispheric coherence plotted over the 1–100 Hz spectrum. Statistical analysis was performed on the average coherence for each frequency band using Two-way ANOVA followed by the Holm-Sidak’s test. (G) Examples of 4-AP-induced epileptic activity in *Chd2* mutant mice. *Chd2*^WT/m^ at 39.4°C, and *Chd2*^m/m^ at 35.4°C. (H) Percentage of mice remaining free of seizures. Statistical analyses: Log-rank test. (I) The proportion of mice with and without seizures (WT: n=6, *Chd2*^WT/m^: n=6, *Chd2*^m/m^: n=4. Statistical analysis: Chi-square test).

Having found increased mRNA levels of *Kcnj11* and *Abcc8* (Fig. 4), we set to examine the physiologic effect of K_ATP_ channels. Interestingly, activating mutations in *KCNJ11* are associated with developmental delay and epilepsy with neonatal or adult-onset diabetes (Ashcroft et al., 2017; Burkart et al., 2024). In some cases, the neurological phenotypes were improved by K_ATP_ inhibitor treatment, such as glibenclamide (GBC) (Slingerland et al., 2006; Mlynarski et al., 2007; Shimomura et al., 2007). The effect of GBC was tested *in vivo*, focusing on the power of background ECoG oscillations. However, acute administration of GBC had no effect on background ECoG oscillations in WT of *Chd2* mutant mice (Supplementary Fig. 6).

Seizures are one of the common phenotypes in *CHD2* patients, and febrile seizures were reported in some *CHD2* patients (Carvill et al., 2015). As the spontaneous epileptic activity was not observed in *Chd2* mutant mice, we tested the sensitivity of *Chd2*^WT/m^ and *Chd2*^m/m^ to heat-induced seizures. However, elevating the body core temperature of these mice up to 42 °C did not cause epileptic activity.

4-aminopyridine (4-AP), a non-selective potassium channel blocker, can have a pro-convulsant effect (Ventura-mejía et al., 2023). Acute administration of 4-AP (4mg/kg) to WT mice did not induce epileptic activity in five out of the six tested mice (83%). Still, it did cause some abnormal nonconvulsive effect in one of the tested mice. Conversely, in *Chd2*^WT/m^ 4-AP induced spontaneous epileptic activity at baseline body core temperature in three of the six tested mice (50%), and epileptic activity after the elevation of body core temperature in two additional mice, causingepileptic activity in 83% of the mice (Fig. 5G-I). In *Chd2*^m/m^ 4-AP triggered spontaneous epileptic activity in 75% of the tested mice (3/4). Spontaneous epileptic activity was observed at baseline temperature in two of the three tested mice (50%), and epileptic activity following temperature elevation in one mouse (25%) (Fig. 5G-I).

Together, quantitative analyses of background ECoG recordings revealed reduced power and coherence in *Chd2* mutant mice, indicating an alteration in brain activity in this model, as well as increased sensitivity to 4-AP-induced epileptic activity.

## Discussion

*CHD2*-DD is characterized by epilepsy, developmental delay, motor deficits, and autistic features (Galizia et al., 2015; Thomas et al., 2015; Lamar and Carvill, 2018; Feng et al., 2022; Prince et al., 2024). Treatment options for this disorder are limited, and there is a need for animal and cellular models to advance the development of new therapeutic avenues.

Here, we used behavioral and electrophysiological analyses to characterize key disease-relevant phenotypes in the juvenile and adult *Chd2* mutant mice, demonstrating alterations in global brain activity, increased sensitivity to epilepsy, motor deficits, and autistic-like features.

There is a complex genotype-phenotype correlation in *CHD2*-related DD and the spectrum of clinical manifestation in patients (Lamar and Carvill, 2018; De Maria et al., 2022; Clara-Hwang et al., 2024; You et al., 2024). Even when focusing only on patients with nonsense and frameshift mutations in the N terminus, which is best mimicked by the model presented here, the clinical presentation varied. The p.E210* variant was detected in a mother with mild controlled epilepsy and high school education. However, in her daughter, the same inherent variant caused refractory epilepsy with global developmental delay and microcephaly (Petersen et al., 2018; You et al., 2024). Similarly, the p.Arg121* variant caused severe epilepsy with developmental delay (Carvill et al., 2013), while the p.S112* variant caused developmental delay without epilepsy (Hamdan et al., 2014). In mice, the strong genetic background effect in *Chd2* mutant mice could be a manifestation of this complexity. Indeed, a modifying effect of genetic background was reported in many other models, including mouse models of absence epilepsy (Frankel, 2009; Frankel et al., 2014), Dravet syndrome (Yu et al., 2006; Hawkins et al., 2016; Calhoun et al., 2017; Mavashov et al., 2023), *SCN2A*-related epileptic encephalopathy (Hawkins and Kearney, 2016), autism (Kazdoba et al., 2016), and a mouse model of *CHD8*-related DD (Tabbaa et al., 2023). Our transcriptomic analyses highlighted *Kcnj11*, a gene associated with developmental delay and epilepsy with neonatal or adult-onset diabetes (Ashcroft et al., 2017; Burkart et al., 2024), as a genetic modifier gene in *Chd2* (Fig. 4).

On the 129X1/SvJ background, *Chd2*^WT/m^ mice, which recapitulate the genetic condition of *CHD2* patients, demonstrated: *i*) reduced CHD2 protein expression; *ii*) reduced power of background ECoG oscillation in multiple frequency bands; *iii*) reduced interhemispheric coherence in the alpha and beta frequency bands; *iv*) increased susceptibility to 4-AP induced epilepsy; *v*) motor deficits; and *vi*) autistic-like features.

In *Chd2*^m/m^, in addition to the above, we observed: *i*) lower levels of CHD2 protein; *ii*) a marked upregulation of the mRNA expression of the mutant allele; *iii*) growth retardation; *iv*) aggravation of some of the motor and nest-building impairment. Nevertheless, we did not detect an exacerbation in the alteration of background ECoG properties or the emergence of spontaneous epileptic activity.

Compared to earlier models, which demonstrated growth retardation (Marfella et al., 2006; Kim et al., 2018; Min et al., 2022), subtle spectral ECoG changes, and impaired long-term spatial memory (Kim et al., 2018), the model presented here provides a more comprehensive representation of the human CHD2-DD phenotypes. First, it highlights the motor deficits and autistic-like features. Second, the clear changes in ECoG power and epileptic susceptibility highlight electrographic disease-relevant biomarkers. Reduced power of background ECoG activity was found to be modulated in multiple neurodevelopmental disorders, including Angelman syndrome, Rett syndrome, and Fragile X (Goodspeed et al., 2023). In Dravet syndrome, low power of background activity correlated with an increased risk of premature death (Fadila et al., 2020; Gerbatin et al., 2022) and was corrected with gene therapy (Fadila et al., 2023). In addition, measurements of seizure susceptibility in Dravet syndrome mice were used to quantify the therapeutic effect of current anti-seizure medications (Hawkins et al., 2017; Pernici et al., 2021; Tanenhaus et al., 2022; Fadila et al., 2023; Quinn et al., 2023). Background power was also proposed as a biomarker in neonatal seizures and hypoxic-ischemic encephalopathy (Korotchikova et al., 2011).

Mechanistically, reduced ECoG power is associated with an imbalance in excitation-inhibition (Gao et al., 2017; Bruining et al., 2020). This is in agreement with the role of CHD2 in interneuron proliferation and migration (Kim et al., 2018; Lamar and Carvill, 2018; Lewis et al., 2022). Thus, these electrographic, quantitative, objective measurements can be a useful tool to examine the therapeutic effect of current and novel treatments for CHD2-DD. Sensitive and quantifiable biomarkers are especially powerful when assessing the therapeutic effect of small molecules, which often lack the ability to completely revert the phenotypes, in contrast to disease-modifying treatments, such as genetic therapies, which often offer correction of the motor and autistic traits (Ozlu et al., 2021; Fadila et al., 2023; Palmieri et al., 2023; Berardino et al., 2024). Here, an examination of GBC’s effect demonstrated that it could not modulate global brain associations in WT or *Chd2* mutants (Supplementary Fig. 6). Still, other potential treatments can be screened in a similar paradigm.

Interestingly, in contrast to previous models of *Chd2* haploinsufficiency, where homozygous knockout mice die perinatally (Marfella et al., 2006; Kulkarni et al., 2008), we did not observe premature mortality in *Chd2*^m/m^. This is consistent with no effects on viability and overall mild phenotypes of *Chd2*^−/−^ mice characterized by the IMPC consortium (Groza et al., 2023). The differences between the models could be related to the low level of residual CHD2 expression in our, and potentially IMPC, mice or to a dominant negative effect of the truncated CHD2 expressed in from the gene trapped allele, which can express a protein from the first 27 exons of CHD2 in the early models (Marfella et al., 2006; Kulkarni et al., 2008). The mechanism for the residual expression in our model is not completely understood and may be due to an alternative initiation site downstream of the frame-shift mutation that can support the production of 1,809 protein, together with a compensatory enhanced transcription (Supplementary Fig. 1)

Overall, the *Chd2* mutant mouse model on the 129X1/SvJ background provides an experimental platform to study the mechanism of CHD2-DD and develop novel and effective treatment options for this intractable disorder.

## Acknowledgments

We acknowledge the financial support of the Coalition to Cure CHD2 (MR), The Israel Science Foundation (#214/22, MR), Recanati Foundation, Faculty of Medical & Health Sciences (MR); Marguerite Stolz Research Fellowship, The Faculty of Medical & Health Sciences (MR), the Abisch-Frenkel RNA Therapeutics Center (IU), the Judith Ruth Institute for Preclinical Brain Research (IU), the Weizmann-Brazil Center for Research on Neurodegeneration (IU), the Gladys Monroy and Larry Marks Center for Brain Disorders (IU), the Nella and Leon Benoziyo Center for Neurological Diseases (IU) and the Prajs-Drimmer Institute, Tel Aviv University (AM).

## Author contributions

Conceptualization: AM, IU and MR; Methodology: AM, ST, YS, MB, RBTP, SQ, YA, KV, IU. MR; Software: SQ; Validation: AM, IU, MR; Formal Analysis: AM, ST, YS, MB, RBTP, SQ, YA, KV, IU. MR; Investigation: AM, ST, YS, MB, RBTP, SQ, YA, KV, IU. MR; Resources: IU, MR; Writing – original draft: AM, MR; Writing – Review & Editing: AM, KV, YS, RBTP, IU, MR; Supervision: IU, MR. Funding Acquisition: IU, MR.

## References

Almog, Y., Fadila, S., Brusel, M., Mavashov, A., Anderson, K., and Rubinstein, M. (2021). Developmental alterations in firing properties of hippocampal CA1 inhibitory and excitatory neurons in a mouse model of Dravet syndrome. Neurobiol. Dis. 148, 105209. doi:10.1016/j.nbd.2020.105209.

Ashcroft, F. M., Puljung, M. C., and Vedovato, N. (2017). Neonatal diabetes and the KATP channel: From mutation to therapy. Trends Endocrinol. Metab. 28, 377–387. doi:10.1016/J.TEM.2017.02.003.

Barak, B., Zhang, Z., Liu, Y., Nir, A., Trangle, S. S., Ennis, M., et al. (2019). Neuronal deletion of Gtf2i, associated with Williams syndrome, causes behavioral and myelin alterations rescuable by a remyelinating drug. Nat. Neurosci. *2019 225* 22, 700–708. doi:10.1038/s41593-019-0380-9.

Barker-Haliski, M., and Hawkins, N. A. (2024). Innovative drug discovery strategies in epilepsy: integrating next-generation syndrome-specific mouse models to address pharmacoresistance and epileptogenesis. Expert Opin. Drug Discov. 19, 1099–1113. doi:10.1080/17460441.2024.2384455.

Bellinvia, A., Portaccio, E., and Amato, M. P. (2023). Current advances in the pharmacological prevention and management of cognitive dysfunction in multiple sclerosis. Expert Opin. Pharmacother. 24, 435–451. doi:10.1080/14656566.2022.2161882.

Berardino, C. Di, Massimino, L., Ungaro, F., and Colasante, G. (2024). Gene therapy for Dravet syndrome: promises and impact on disease trigger and secondary modifications. Rare Dis Orphan Drugs J *2024;321*. 3, N/A-N/A. doi:10.20517/RDODJ.2024.07.

Bowyer, S. M. (2016). Coherence a measure of the brain networks: past and present. Neuropsychiatr. Electrophysiol. 2, 1–12. doi:10.1186/s40810-015-0015-7.

Bruining, H., Hardstone, R., Juarez-Martinez, E. L., Sprengers, J., Avramiea, A. E., Simpraga, S., et al. (2020). Measurement of excitation-inhibition ratio in autism spectrum disorder using critical brain dynamics. Sci. Rep. 10. doi:10.1038/S41598-020-65500-4.

Burkart, M.-E., Kurzke, J., Jacobi, R., Vera, J., Ashcroft, F. M., Eilers, J., et al. (2024). KATP channel mutation disrupts hippocampal network activity and nocturnal gamma shifts. Brain. doi:10.1093/BRAIN/AWAE157.

Calhoun, J. D., Hawkins, N. A., Zachwieja, N. J., and Kearney, J. A. (2017). Cacna1g is a genetic modifier of epilepsy in a mouse model of Dravet syndrome. Epilepsia 58, e111– e115. doi:10.1111/EPI.13811.

Carvill, G., Helbig, I., and Mefford, H. (2015). CHD2-related neurodevelopmental disorders. GeneReviews®.

Carvill, G. L., Heavin, S. B., Yendle, S. C., McMahon, J. M., O’Roak, B. J., Cook, J., et al. (2013). Targeted resequencing in epileptic encephalopathies identifies de novo mutations in CHD2 and SYNGAP1. Nat. Genet. 45, 825–830. doi:10.1038/ng.2646.

Clara-Hwang, A., Stefani, S., Lau, T., Scala, M., Aynekin, B., Bernardo, P., et al. (2024). Expanding the mutational landscape and clinical phenotype of CHD2-related encephalopathy. Neurol. Genet. 10. doi:10.1212/NXG.0000000000200168.

Cook, A. A., Fields, E., and Watt, A. J. (2021). Losing the Beat: Contribution of Purkinje Cell Firing Dysfunction to Disease, and Its Reversal. Neuroscience 462, 247–261. doi:10.1016/J.NEUROSCIENCE.2020.06.008.

De Maria, B., Balestrini, S., Mei, D., Melani, F., Pellacani, S., Pisano, T., et al. (2022). Expanding the genetic and phenotypic spectrum of CHD2-related disease: From early neurodevelopmental disorders to adult-onset epilepsy. Am. J. Med. Genet. Part A 188, 522–533. doi:10.1002/AJMG.A.62548.

Deacon, R. M. J. (2006). Assessing nest building in mice. Nat. Protoc. 1, 1117–1119. doi:10.1038/nprot.2006.170.

Dietrich, M., Hartung, H. P., and Albrecht, P. (2021). Neuroprotective Properties of 4-Aminopyridine. Neurol. Neuroimmunol. neuroinflammation 8. doi:10.1212/NXI.0000000000000976.

Dobin, A., Davis, C. A., Schlesinger, F., Drenkow, J., Zaleski, C., Jha, S., et al. (2013). STAR: ultrafast universal RNA-seq aligner. Bioinformatics 29, 15–21. doi:10.1093/BIOINFORMATICS/BTS635.

Dutta, S., and Sengupta, P. (2016). Men and mice: Relating their ages. Life Sci. 152, 244–248. doi:10.1016/J.LFS.2015.10.025.

Fadila, S., Beucher, B., Dopeso-reyes, I. G., Mavashov, A., Brusel, M., Anderson, K., et al. (2023). Viral vector–mediated expression of NaV1.1, after seizure onset, reduces epilepsy in mice with Dravet syndrome. J. Clin. Invest. 133, e159316.

Fadila, S., Quinn, S., Turchetti Maia, A., Yakubovich, D., Ovadia, M., Anderson, K. L., et al. (2020). Convulsive seizures and some behavioral comorbidities are uncoupled in the Scn1aA1783V Dravet syndrome mouse model. Epilepsia 61, 2289–2300. doi:10.1111/epi.16662.

Feng, W., Fang, F., Wang, X., Chen, C., Lu, J., and Deng, J. (2022). Clinical analysis of CHD2 gene mutations in pediatric patients with epilepsy. Pediatr. Investig. 6, 93–99. doi:10.1002/PED4.12321.

Frankel, W. N. (2009). Genetics of complex neurological disease: challenges and opportunities for modeling epilepsy in mice and rats. Trends Genet. 25, 361–367. doi:10.1016/j.tig.2009.07.001.

Frankel, W. N., Mahaffey, C. L., McGarr, T. C., Beyer, B. J., and Letts, V. A. (2014). Unraveling Genetic Modifiers in the Gria4 Mouse Model of Absence Epilepsy. PLOS Genet. 10, e1004454. doi:10.1371/JOURNAL.PGEN.1004454.

Galizia, E. C., Myers, C. T., Leu, C., De Kovel, C. G. F., Afrikanova, T., Cordero-Maldonado, M. L., et al. (2015). CHD2 variants are a risk factor for photosensitivity in epilepsy. Brain 138, 1198–1208. doi:10.1093/BRAIN/AWV052.

Gao, R., Peterson, E. J., and Voytek, B. (2017). Inferring synaptic excitation/inhibition balance from field potentials. Neuroimage 158, 70–78. doi:10.1016/j.neuroimage.2017.06.078.

Gerbatin, R. R., Augusto, J., Boutouil, H., Reschke, C. R., and Henshall, D. C. (2022). Life-span characterization of epilepsy and comorbidities in Dravet syndrome mice carrying a targeted deletion of exon 1 of the Scn1a gene. Exp. Neurol. 354, 114090. doi:10.1016/J.EXPNEUROL.2022.114090.

Goodspeed, K., Armstrong, D., Dolce, A., Evans, P., Said, R., Tsai, P., et al. (2023). Electroencephalographic (EEG) Biomarkers in Genetic Neurodevelopmental Disorders. J. Child Neurol. 38, 466. doi:10.1177/08830738231177386.

Groza, T., Gomez Lopez, F. L., Mashhadi, H. H., Muñoz-Fuentes, V., Gunes, O., Wilson, R., et al. (2023). The International Mouse Phenotyping Consortium: comprehensive knockout phenotyping underpinning the study of human disease. Nucleic Acids Res. 51, D1038–D1045. doi:10.1093/NAR/GKAC972.

Hamdan, F. F., Srour, M., Capo-Chichi, J. M., Daoud, H., Nassif, C., Patry, L., et al. (2014). De novo mutations in moderate or severe intellectual disability. PLoS Genet. 10, e1004772. doi:10.1371/journal.pgen.1004772.

Hawkins, N. A., Anderson, L. L., Gertler, T. S., Laux, L., George, A. L., and Kearney, J. A. (2017). Screening of conventional anticonvulsants in a genetic mouse model of epilepsy. Ann. Clin. Transl. Neurol. 4, 326–339. doi:10.1002/acn3.413.

Hawkins, N. A., and Kearney, J. A. (2016). Hlf is a genetic modifier of epilepsy caused by voltage-gated sodium channel mutations. Epilepsy Res. 119, 20–23. doi:10.1016/J.EPLEPSYRES.2015.11.016.

Hawkins, N. A., Zachwieja, N. J., Miller, A. R., Anderson, L. L., and Kearney, J. A. (2016). Fine Mapping of a Dravet Syndrome Modifier Locus on Mouse Chromosome 5 and Candidate Gene Analysis by RNA-Seq. PLoS Genet. 12, e1006398. doi:10.1371/journal.pgen.1006398.

Hedrich, U. B. S., Lauxmann, S., Wolff, M., Synofzik, M., Bast, T., Binelli, A., et al. (2021). 4-Aminopyridine is a promising treatment option for patients with gain-of-function KCNA2-encephalopathy. Sci. Transl. Med. 13. doi:10.1126/SCITRANSLMED.AAZ4957.

Jirkof, P. (2014). Burrowing and nest building behavior as indicators of well-being in mice. J. Neurosci. Methods 234, 139–146. doi:10.1016/J.JNEUMETH.2014.02.001.

Kazdoba, T. M., Leach, P. T., and Crawley, J. N. (2016). Behavioral phenotypes of genetic mouse models of autism. *Genes*, Brain Behav. 15, 7–26. doi:10.1111/gbb.12256.

Kim, Y. J., Khoshkhoo, S., Frankowski, J. C., Zhu, B., Abbasi, S., Lee, S., et al. (2018). Chd2 is necessary for neural circuit development and long-term memory. Neuron 100, 1180–1193.e6. doi:10.1016/j.neuron.2018.09.049.

Korotchikova, I., Stevenson, N. J., Walsh, B. H., Murray, D. M., and Boylan, G. B. (2011). Quantitative EEG analysis in neonatal hypoxic ischaemic encephalopathy. Clin. Neurophysiol. 122, 1671–1678. doi:10.1016/j.clinph.2010.12.059.

Kulkarni, S., Nagarajan, P., Wall, J., Donovan, D. J., Donell, R. L., Ligon, A. H., et al. (2008). Disruption of chromodomain helicase DNA binding protein 2 (CHD2) causes scoliosis. Am. J. Med. Genet. Part A 146A, 1117–1127. doi:10.1002/AJMG.A.32178.

Lamar, K. M. J., and Carvill, G. L. (2018). Chromatin remodeling proteins in epilepsy: Lessons from CHD2-associated epilepsy. Front. Mol. Neurosci. 11, 208. doi:10.3389/fnmol.2018.00208.

Lewis, E. M. A., Chapman, G., Kaushik, K., Determan, J., Antony, I., Meganathan, K., et al. (2022). Regulation of human cortical interneuron development by the chromatin remodeling protein CHD2. Sci. Rep. 12, 1–18. doi:10.1038/s41598-022-19654-y.

Li, B., and Dewey, C. N. (2011). RSEM: accurate transcript quantification from RNA-Seq data with or without a reference genome. BMC Bioinformatics 12. doi:10.1186/1471-2105-12-323.

Love, M. I., Huber, W., and Anders, S. (2014). Moderated estimation of fold change and dispersion for RNA-seq data with DESeq2. Genome Biol. 15. doi:10.1186/S13059-014-0550-8.

Malone, H. A., and Roberts, C. W. M. (2024). Chromatin remodellers as therapeutic targets. Nat. Rev. Drug Discov. 23, 661–681. doi:10.1038/s41573-024-00978-5.

Marfella, C. G. A., Ohkawa, Y., Coles, A. H., Garlick, D. S., Jones, S. N., and Imbalzano, A. N. (2006). Mutation of the SNF2 family member Chd2 affects mouse development and survival. J. Cell. Physiol. 209, 162–171. doi:10.1002/JCP.20718.

Mavashov, A., Brusel, M., Liu, J., Woytowicz, V., Bae, H., Chen, Y.-H., et al. (2023). Heat-induced seizures, premature mortality, and hyperactivity in a novel *Scn1a* nonsense model for Dravet syndrome. Front. Cell. Neurosci. 17, 1149391. doi:10.3389/FNCEL.2023.1149391.

McGraw, C. M., Ward, C. S., and Samaco, R. C. (2017). Genetic rodent models of brain disorders: Perspectives on experimental approaches and therapeutic strategies. Am. J. Med. Genet. C. Semin. Med. Genet. 175, 368. doi:10.1002/AJMG.C.31570.

Min, Z., Xin, H., Liu, X., Wan, J., Fan, Z., Rao, X., et al. (2022). Chromodomain helicase DNA-binding domain 2 maintains spermatogonial self-renewal by promoting chromatin accessibility and mRNA stability. iScience 25, 105552. doi:10.1016/J.ISCI.2022.105552.

Mlynarski, W., Tarasov, A. I., Gach, A., Girard, C. A., Pietrzak, I., Zubcevic, L., et al. (2007). Sulfonylurea improves CNS function in a case of intermediate DEND syndrome caused by a mutation in KCNJ11. Nat. Clin. Pract. Neurol. 3, 640–645. doi:10.1038/NCPNEURO0640.

Ozlu, C., Bailey, R. M., Sinnett, S., and Goodspeed, K. D. (2021). Gene transfer therapy for neurodevelopmental disorders. Dev. Neurosci. 43, 230–240. doi:10.1159/000515434.

Palmieri, M., Pozzer, D., and Landsberger, N. (2023). Advanced genetic therapies for the treatment of Rett syndrome: state of the art and future perspectives. Front. Neurosci. 17, 1172805. doi:10.3389/FNINS.2023.1172805/BIBTEX.

Pernici, C. D., Mensah, J. A., Dahle, E. J., Johnson, K. J., Handy, L., Buxton, L., et al. (2021). Development of an antiseizure drug screening platform for Dravet syndrome at the NINDS contract site for the Epilepsy Therapy Screening Program. Epilepsia 62, 1665–1676. doi:10.1111/epi.16925.

Petersen, A. K., Streff, H., Tokita, M., and Bostwick, B. L. (2018). The first reported case of an inherited pathogenic CHD2 variant in a clinically affected mother and daughter. Am. J. Med. Genet. A 176, 1667–1669. doi:10.1002/AJMG.A.38835.

Prince, S., Bonkowski, E., McGraw, C., SanInocencio, C., Mefford, H. C., Carvill, G., et al. (2024). A roadmap to cure CHD2-related disorders. Ther. Adv. Rare Dis. 5, 26330040241283748. doi:10.1177/26330040241283749.

Quinn, S., Brusel, M., Ovadia, M., and Rubinstein, M. (2023). Acute effect of antiseizure drugs on background oscillations in Scn1aA1783V Dravet syndrome mouse model. Front. Pharmacol. 14, 703. doi:10.3389/FPHAR.2023.1118216.

Rom, A., Melamed, L., Gil, N., Goldrich, M. J., Kadir, R., Golan, M., et al. (2019). Regulation of CHD2 expression by the Chaserr long noncoding RNA gene is essential for viability. Nat. Commun. 10. doi:10.1038/s41467-019-13075-8.

Satterstrom, F. K., Kosmicki, J. A., Wang, J., Breen, M. S., De Rubeis, S., An, J. Y., et al. (2020). Large-scale exome sequencing study implicates both developmental and functional changes in the neurobiology of autism. Cell 180, 568–584.e23. doi:10.1016/j.cell.2019.12.036.

Shevelkin, A. V., Ihenatu, C., and Pletnikov, M. V. (2014). Pre-clinical models of neurodevelopmental disorders: focus on the cerebellum. Rev. Neurosci. 25, 177. doi:10.1515/REVNEURO-2013-0049.

Shimomura, K., Hörster, F., De Wet, H., Flanagan, S. E., Ellard, S., Hattersley, A. T., et al. (2007). A novel mutation causing DEND syndrome: a treatable channelopathy of pancreas and brain. Neurology 69, 1342–1349. doi:10.1212/01.WNL.0000268488.51776.53.

Silverman, J. L., Yang, M., Lord, C., and Crawley, J. N. (2010). Behavioural phenotyping assays for mouse models of autism. Nat. Rev. Neurosci. *2010 117* 11, 490–502. doi:10.1038/nrn2851.

Slingerland, A. S., Nuboer, R., Hadders-Algra, M., Hattersley, A. T., and Bruining, G. J. (2006). Improved motor development and good long-term glycaemic control with sulfonylurea treatment in a patient with the syndrome of intermediate developmental delay, early-onset generalised epilepsy and neonatal diabetes associated with the V59M mutation in the KCNJ11 gene. Diabetologia 49, 2559–2563. doi:10.1007/S00125-006-0407-0.

Suls, A., Jaehn, J. A., Kecskés, A., Weber, Y., Weckhuysen, S., Craiu, D. C., et al. (2013). De novo loss-of-function mutations in CHD2 cause a fever-sensitive myoclonic epileptic encephalopathy sharing features with Dravet syndrome. Am. J. Hum. Genet. 93, 967– 975. doi:10.1016/j.ajhg.2013.09.017.

Tabbaa, M., Knoll, A., and Levitt, P. (2023). Mouse population genetics phenocopies heterogeneity of human Chd8 haploinsufficiency. Neuron 111, 539–556.e5. doi:10.1016/J.NEURON.2023.01.009.

Tanenhaus, A., Stowe, T., Young, A., McLaughlin, J., Aeran, R., Lin, I. W., et al. (2022). Cell-selective adeno-associated virus-mediated SCN1A gene regulation therapy rescues mortality and seizure phenotypes in a Dravet syndrome mouse model and is well tolerated in nonhuman primates. Hum. Gene Ther. 33, 579–597. doi:10.1089/hum.2022.037.

Tang, X., and Sanford, L. D. (2005). Home cage activity and activity-based measures of anxiety in 129P3/J, 129X1/SvJ and C57BL/6J mice. Physiol. Behav. 84, 105–115. doi:10.1016/J.PHYSBEH.2004.10.017.

Thomas, R. H., Zhang, L. M., Carvill, G. L., Archer, J. S., Heavin, S. B., Mandelstam, S. A., et al. (2015). CHD2 myoclonic encephalopathy is frequently associated with self-induced seizures. Neurology 84, 951. doi:10.1212/WNL.0000000000001305.

Ventura-mejía, C., Nuñez-ibarra, B. H., and Medina-ceja, L. (2023). An update of 4-aminopyride as a useful model of generalized seizures for testing antiseizure drugs: in vitro and in vivo studies. Acta Neurobiol. Exp. (Wars). 83, 63–70. doi:10.55782/ane-2023-007.

Wang, W., and Frankel, W. N. (2021). Overlaps, gaps, and complexities of mouse models of Developmental and Epileptic Encephalopathy. Neurobiol. Dis. 148, 105220. doi:10.1016/J.NBD.2020.105220.

Woo, S. K., Tsymbalyuk, N., Tsymbalyuk, O., Ivanova, S., Gerzanich, V., and Simard, J. M. (2020). SUR1-TRPM4 channels, not KATP, mediate brain swelling following cerebral ischemia. Neurosci. Lett. 718. doi:10.1016/J.NEULET.2019.134729.

You, C., Xu, L., Zhu, L., Qiu, S., Xu, N., Wang, Y., et al. (2024). Clinical analysis of five CHD2 gene mutations in Chinese children with epilepsy. Seizure 121, 38–44. doi:10.1016/j.seizure.2024.07.009.

Yu, F. H., Mantegazza, M., Westenbroek, R. E., Robbins, C. A., Kalume, F., Burton, K. A., et al. (2006). Reduced sodium current in GABAergic interneurons in a mouse model of severe myoclonic epilepsy in infancy. Nat. Neurosci. 9, 1142–1149. doi:10.1038/nn1754.

Zhu, L., Peng, F., Deng, Z., Feng, Z., and Ma, X. (2022). A novel variant of the CHD2 gene associated with developmental delay and myoclonic epilepsy. Front. Genet. 13, 761178. doi:10.3389/FGENE.2022.761178/BIBTEX.

