## Supplementary Material for "Alterations in background ECoG activity and behavioral deficits in a mouse model of CHD2-related developmental delay"

Supplementary Figs. 1-6
Supplementary Table 1

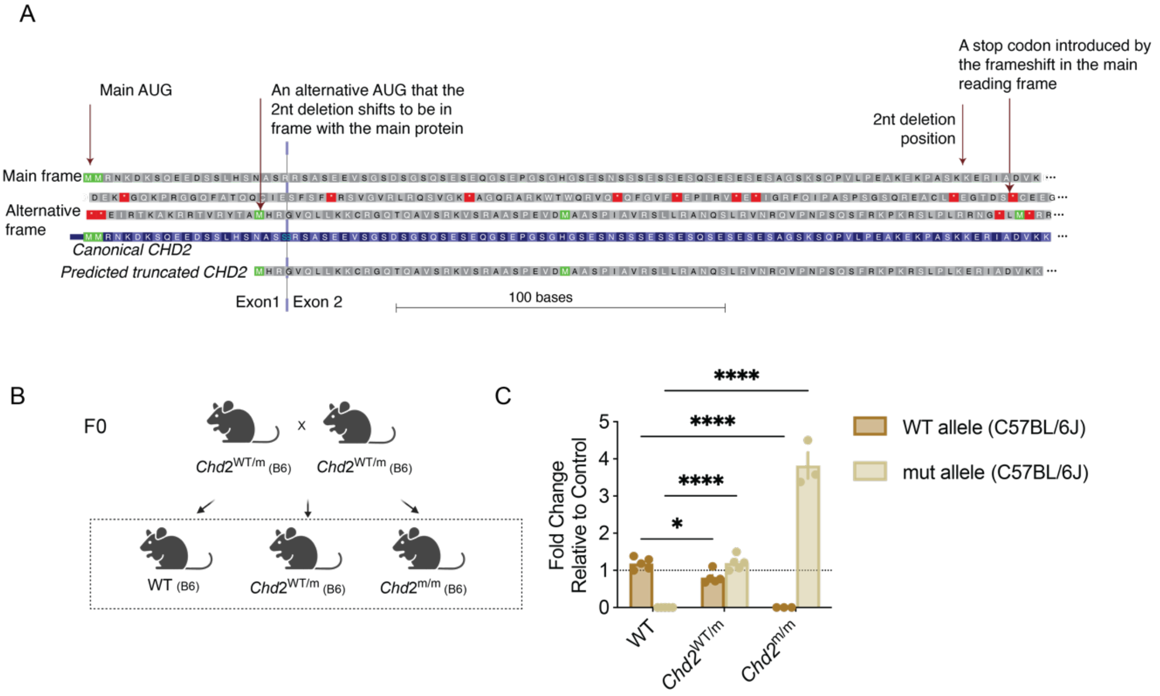


Supplementary Fig. 1. **CHD2 sequence in WT and *Chd2* mutant mice.**

(A) CRISPR-mediated deletion of nucleotides 266 and 267 in the *Chd2* caused a frameshift after a.a 89, changing the canonical sequence of KERI to GTDS*. The same frameshift also resulted in the creation of an alternative start codon that the mutation puts back into the frame with most of the protein (bottom, predicted truncated CHD2).

(B) Depiction of the breeding scheme to generate WTs, *Chd2*$^{WT/m}$, and *Chd2*$^{m/m}$ on the pure C57BL/6J (B6) background.

(C) Allele-specific quantification of *Chd2* transcripts in mice on the pure C57BL/6J background (WT: n=5, *Chd2*$^{WT/m}$: n=5, *Chd2*$^{m/m}$: n=3. Statistical analysis: Two-way ANOVA followed by Holm-Sidak's test).

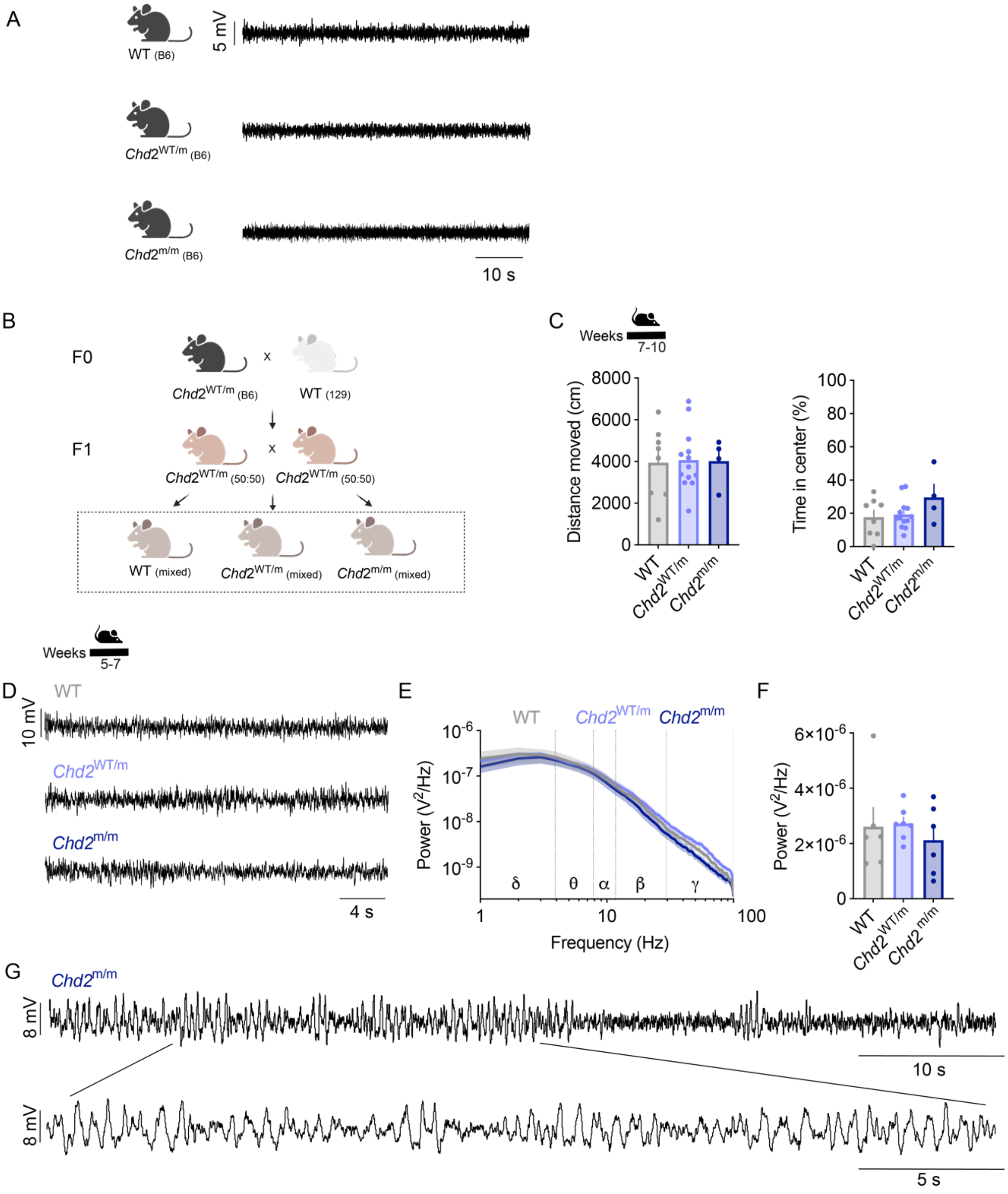


Supplementary Fig. 2. ***Chd2* mutant mice on the C57BL/6J background or on a mixed B6:129 genetic background do not show clear neurological phenotypes.**

(A) Example traces of background ECoG activity of WT, *Chd2*$^{WT/m}$, and *Chd2*$^{m/m}$ on a pure C57BL/6J background.
(B) Depiction of the breeding scheme. *Chd2*$^{WT/m}$ B6 F0 mice were crossbred with 129X1/SvJ WT mice to generate *Chd2*$^{WT/m}$ mice on a 50:50 B6:129 mixed genetic background. The progeny of mice on a 50:50 genetic background is depicted in this Figure (marked by the dashed line). (C) The exploration of adult mice (7-10 weeks old) was tested in the open field. The total distance traveled and the time in the center are depicted (WT: n=8 (F=3, M=5), *Chd2*$^{WT/m}$: n=13 (F=6, M=7), *Chd2*$^{m/m}$: n=4 (F=2, M=2). Statistical analysis: One-way ANOVA followed by Dunnett's test). (D-G) ECoG recordings. (D) Examples of background ECoG activity. (E) Power spectral density. (F) Total power. (WT: n=6 (F=3, M=3), *Chd2*$^{WT/m}$: n=7 (F=3, M=4), *Chd2*$^{m/m}$: n=6 (F=3, M=3). Statistical analysis: Kruskal-Wallis test followed by Dunn's test. The recorded mice were 5-7 weeks old. (G) Nonconvulsive epileptic activity was detected in one *Chd2*$^{m/m}$ recorded at the age of five weeks.

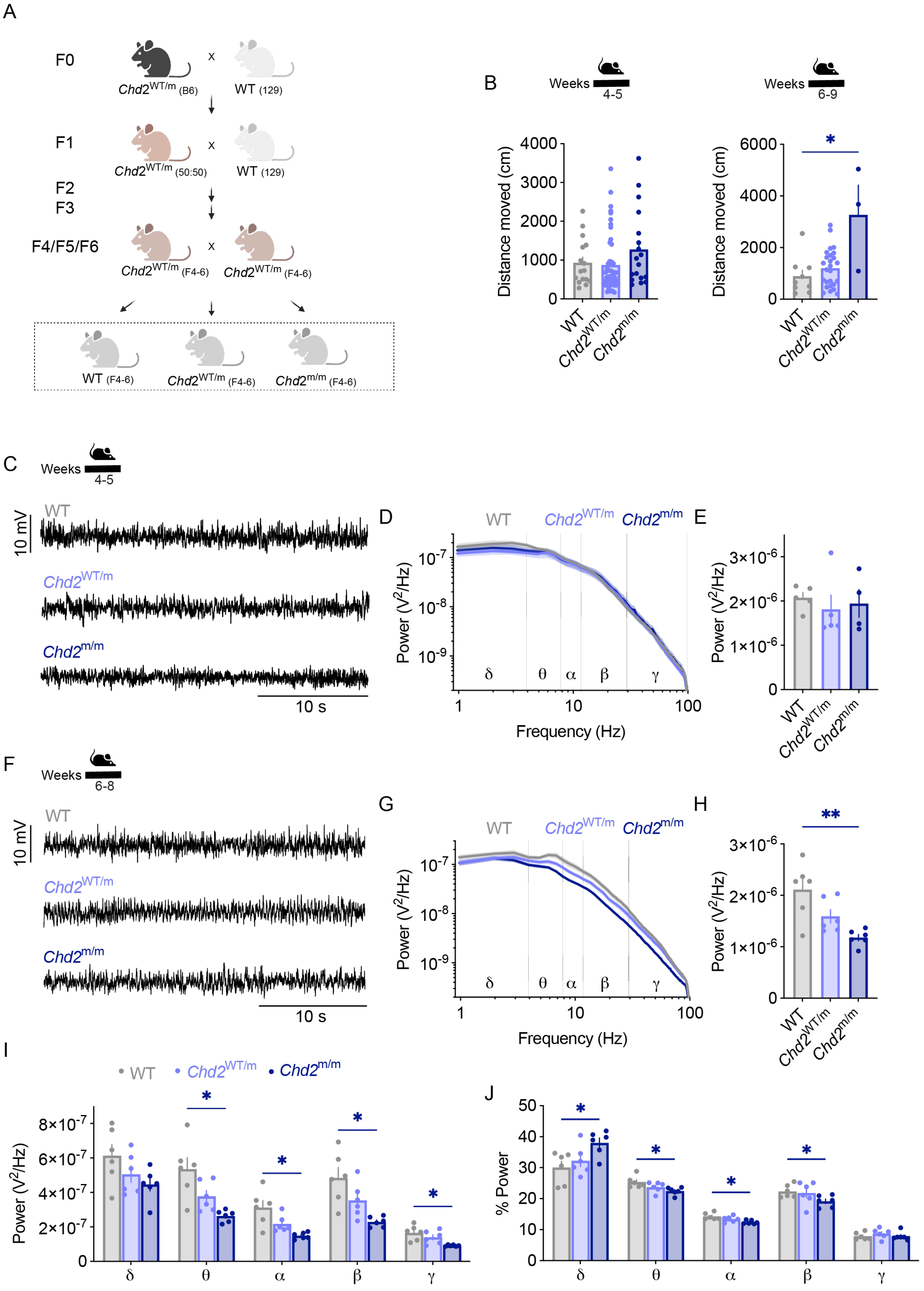


Supplementary Fig. 3. **Reduced ECoG power in *Chd2*$^{m/m}$ mice on an enriched 129x1/SvJ background.** (A) Depiction of the breeding scheme. *Chd2*$^{WT/m}$ mice were crossed with WT mice on the pure 129x1/SvJ for at least four generations prior to the breeding of the two *Chd2*$^{WT/m}$ mice on an enriched 129 background together. The dashed line marks mice that were used to generate the depicted data. (B) The total distance moved in the open field of mice at the age of 4-5 weeks (left: WT: n=16 (F=9, M=7), *Chd2*$^{WT/m}$: n=49 (F=30 ,

M=19), *Chd2*$^{m/m}$: n=18 (F=15, M=3). Statistical analysis: Kruskal-Wallis test followed by Dunn's test) or 6-9 weeks (right: WT: n=9 (F=5, M=4), *Chd2*$^{WT/m}$: n=26 (F=13, M=13), *Chd2*$^{m/m}$: n=3 (F=3). Statistical analysis: Kruskal-Wallis test followed by Dunn's test).
(C-E) ECoG recordings in mice at the age of 4-5 weeks. (C) Examples of background ECoG activity. (D) Power spectral density. (E) Total power. (WT: n=5 (F=3, M=2), *Chd2*$^{WT/m}$: n=5 (F=3, M=2), *Chd2*$^{m/m}$: n=4 (F=3, M=1). Statistical analysis: Kruskal-Wallis test followed by Dunn's test).
(F-I) ECoG recordings in mice at the age of 6-8 weeks. (F) Examples of background ECoG activity. (G) Power spectral density. (H) Total power. (WT: n=6 (F=3, M=3), *Chd2*$^{WT/m}$: n=6 (F=3, M=3), *Chd2*$^{m/m}$: n=6 (F=4, M=2). Statistical analysis: One-way ANOVA followed by Dunnett's test). (I) Total power in the following frequency bands: δ, 0.9–3.9 Hz; θ, 4.8–7.8 Hz; α, 8.7–11.7 Hz; β, 12.6–29.2 Hz; γ, 30.2–99.6 Hz. Statistical analysis: Two-way ANOVA followed by the Holm-Sidak's test. (J) The normalized power (Sum of the power in each band/total power) in each frequency band. Statistical analysis: Two-way ANOVA followed by the Holm-Sidak's test.

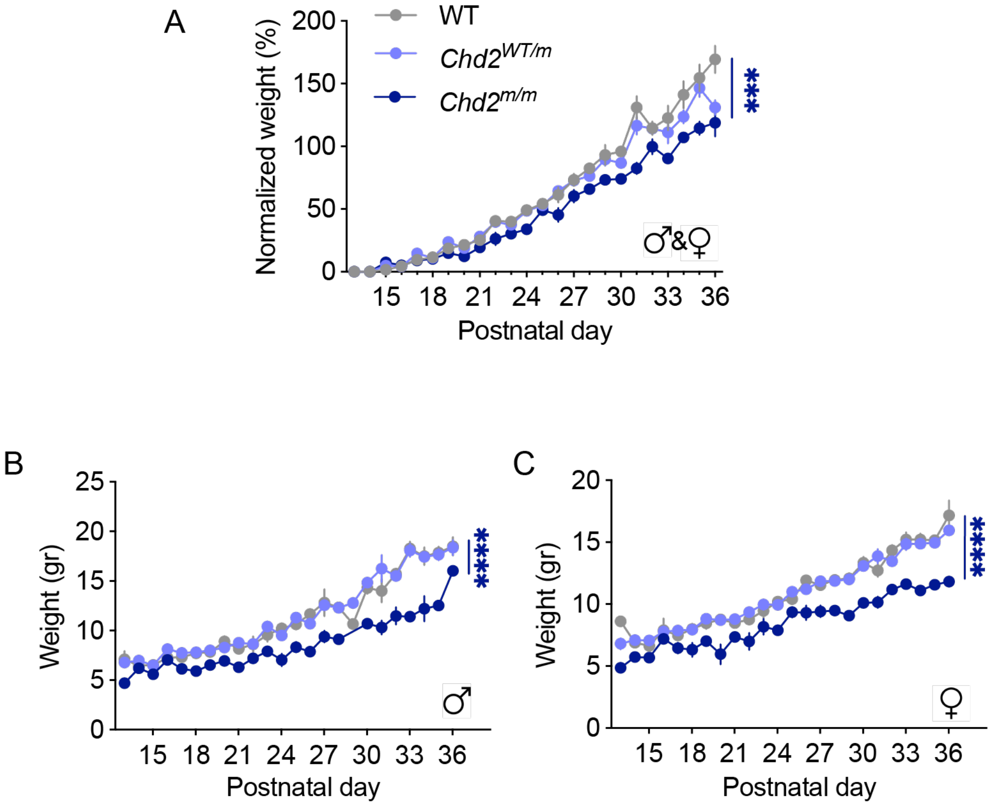


Supplementary Fig. 4. **Reduced growth in *Chd2*$^{m/m}$ mice**.
(A) Normalized weight ((Δgained weight/initial weight)x100). (B-C) Reduced weight over development in males (B) and females (C) (WT: n=40, (F=20, M=20); *Chd2*$^{WT/m}$: n=64 (F=36, m=28); *Chd2*$^{m/m}$: n=34 (F=17, M=17). Statistical analysis: Mixed model ANOVA followed by Tukey's test).

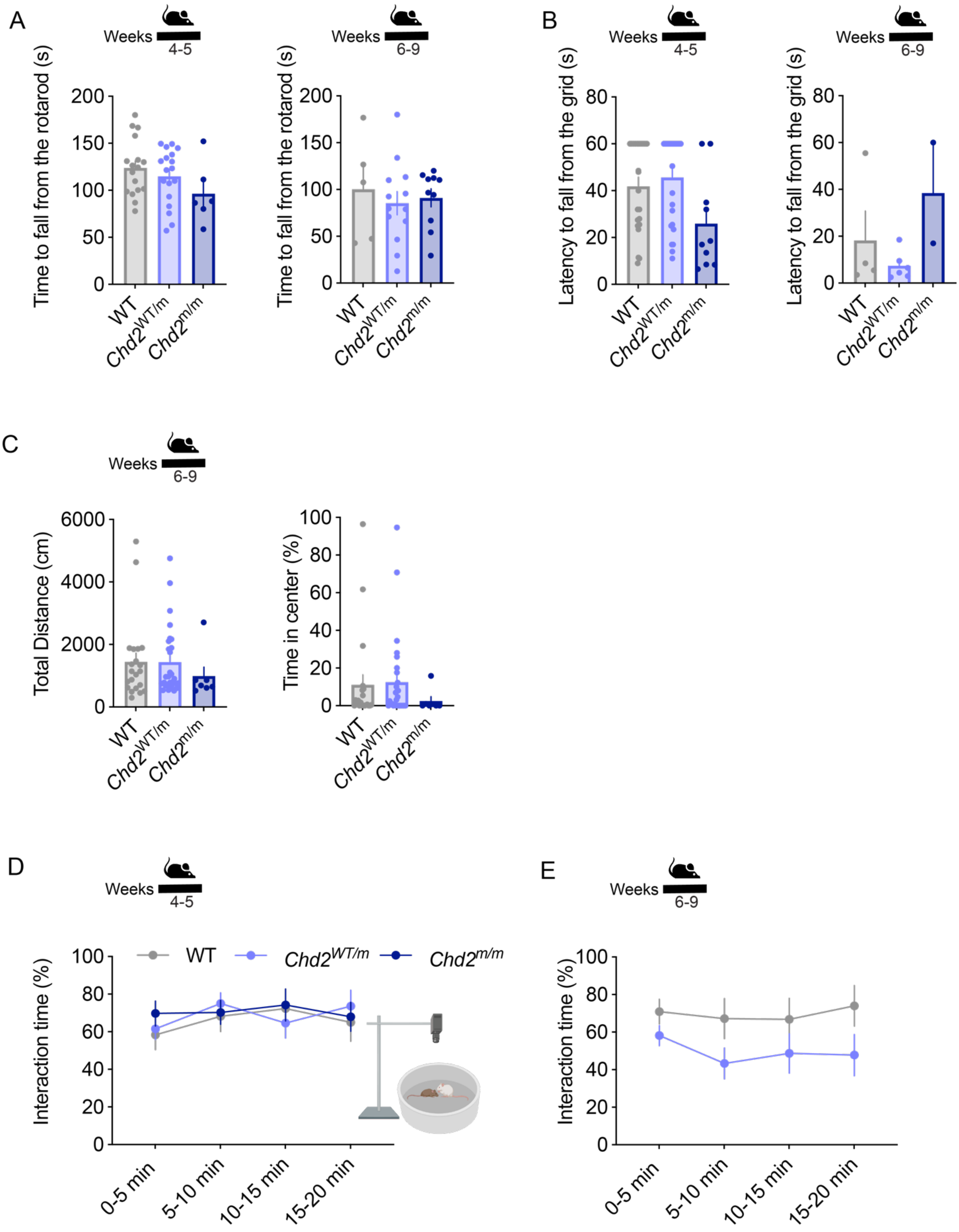


Supplementary Fig. 5. **Additional behavioral data in mice that were crossed onto the 129X1/SvJ background for at least seven generations.**
(A) The latency to fall from the rotarod in juvenile mice (left, WT: n=17 (F=7, M=10), *Chd2*$^{WT/m}$: n=18 (F=9, M=9), *Chd2*$^{m/m}$: n=6 (F=3, M=3). Statistical analysis: One-way ANOVA followed by Dunnett's test) and adult mice (right, WT: n=5 (F=1, M=4), *Chd2*$^{WT/m}$: n=13 (F=6, M=6), *Chd2*$^{m/m}$: n=10 (F=6, M=4). Statistical analysis: One-way ANOVA followed by Dunnett's test). (B) The latency to fall from an up-side-down grid in the four limb-hanging tests in juvenile mice (left, WT: n=23 (F=6. M=17), *Chd2*$^{WT/m}$: n=26 (F=11, M=15), *Chd2*$^{m/m}$: n=10 (F=4, M=6). Statistical analysis: Kruskal-Wallis test followed by Dunn's test) and adult mice (right, WT: n=4 (F=2, M=2), *Chd2*$^{WT/m}$: n=6 (F=6), *Chd2*$^{m/m}$: n=2 (F=1, M=1). Statistical analysis: Kruskal-Wallis test followed by Dunn's test). (C) The total distance traveled in the open field (left) and the percent of time spent in the

center of the arena (right) (Adult mice. WT: n=21 (F=9, M=12), *Chd2*$^{WT/m}$: n=28 (F=14, M=14), *Chd2*$^{m/m}$: n=7 (M=7). Statistical analysis: Kruskal-Wallis test followed by Dunn's test). (D) Social interaction time between juvenile mice. A WT interacting mouse and a test mouse (WT, *Chd2*$^{WT/m}$ or *Chd2*$^{m/m}$) were placed in an arena; their interaction time is depicted in 5 min bins (WT: n=13 (F=5, M=8), *Chd2*$^{WT/m}$: n=14 (F=7, M=7), *Chd2*$^{m/m}$: n=13 (F=8, M=5). Statistical analysis: Two-way ANOVA followed by the Holm-Sidak's test). (E) Social interaction time between adult mice. A WT interacting mouse and a test mouse (WT or *Chd2*$^{WT/m}$) were placed in an arena; their interaction time is depicted in 5 min bins. (WT: n=9 (F=5, M=4), *Chd2*$^{WT/m}$: n=12 (F=8, M=4). Statistical analysis: Two-way ANOVA followed by the Holm-Sidak's test).

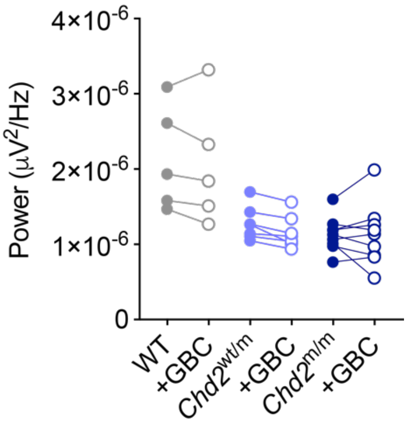


Supplementary Fig. 6. **Glibenclamide (1 mg/kg) did not affect the power of background oscillations.** The mice were recorded in two consecutive sections, two hours before and two hours after an intraperitoneal injection of 1 mg/kg GBC. The power of WT and *Chd2* mutant mice was calculated over the whole section, excluding the first 30 min in each section (the time needed for habituation to the recording chamber and drug absorption) (WT: n=5 (F=2, M=3), *Chd2*$^{WT/m}$: n=7 (F=1, F=6), *Chd2*$^{m/m}$: n=9 (F=5, M=4). Statistical analysis: Two-Way ANOVA followed by Tukey's test).

**Supplementary Table 1: Full statistical information**

| Figure | | Test used | The number of mice | P value | Post-hoc analysis |
|---|---|---|---|---|---|
| Fig. 1 | D | One-way ANOVA | WT: n=3<br>*Chd2*$^{WT/m}$: n=3<br>*Chd2*$^{m/m}$: n=3 | p=0.036 | Holm-Sidak's:<br>p=0.0357 for WT vs. *Chd2*$^{WT/m}$<br>p=0.034 for WT vs. *Chd2*$^{m/m}$ |
| | E | Two-Way ANOVA | WT: n=3<br>*Chd2*$^{WT/m}$: n=3<br>*Chd2*$^{m/m}$: n=3 | p=0.0018 for the main effect of genotype<br>p=0.0012 for the main effect of allele | Dunnett's test:<br>WT allele:<br>p=0.43 for WT vs. *Chd2*$^{WT/m}$<br>p=0.0462 for WT vs. *Chd2*$^{m/m}$<br>Mut allele:<br>p=0.0369 for WT vs. *Chd2*$^{WT/m}$<br>p<0.0001 for WT vs. *Chd2*$^{m/m}$ |
| | G | Mixed-model ANOVA | WT: n=40 (F=20, M=20)<br>*Chd2*$^{WT/m}$: n=64 (F=36, M=28)<br>*Chd2*$^{m/m}$: n=34 (F=17, M=17) | p<0.0001 for the main effect of genotype | Tukey's test:<br>p=0.92 for WT vs. *Chd2*$^{WT/m}$<br>p<0.0001 for WT vs. *Chd2*$^{m/m}$ |
| | H | Mixed-model ANOVA | WT: n=40 (F=20, M=20)<br>*Chd2*$^{WT/m}$: n=64 (F=36, M=28)<br>*Chd2*$^{m/m}$: n=34 (F=17, M=17) | p<0.0001 for the main effect of genotype | Tukey's test:<br>p=0.86 for WT vs. *Chd2*$^{WT/m}$<br>p<0.0001 for WT vs. *Chd2*$^{m/m}$ |
| | J | One-Way ANOVA | WT: n=7 (F=3, M=4)<br>*Chd2*$^{WT/m}$: n=7 (F=2, M=5)<br>*Chd2*$^{m/m}$: n=7 (F=3, M=4) | p=0.28 | Holm-Sidak's:<br>p=0.99 for WT vs. *Chd2*$^{WT/m}$<br>p=0.312 for WT vs. *Chd2*$^{m/m}$ |
| Fig. 2 | B | Kruskal-Wallis | Left:<br>WT: n=22 (F=10, M=12)<br>*Chd2*$^{WT/m}$: n=34 (F=25, M=9)<br>*Chd2*$^{m/m}$: n=10 (F=7, M=3)<br>Right:<br>WT: n=7 (F=4, M=3)<br>*Chd2*$^{WT/m}$: n=14 (F=6, M=8)<br>*Chd2*$^{m/m}$: n=6 (F=5, M=1) | Left:<br>p<0.0001<br>Right:<br>p=0.0002 | Dunn's test:<br>Left:<br>p=0.031 for WT vs. *Chd2*$^{WT/m}$<br>p<0.0001 for WT vs. *Chd2*$^{m/m}$<br>Right:<br>p=0.204 for WT vs. *Chd2*$^{WT/m}$<br>p=0.0001 for WT vs. *Chd2*$^{m/m}$ |
| | C | Kruskal-Wallis | WT: n=23 (F=11, M=12)<br>*Chd2*$^{WT/m}$: n=28 (F=11, M=17)<br>*Chd2*$^{m/m}$: n=10 (F=5, M=5) | Left:<br>p=0.64<br>Right:<br>p=0.0129 | Dunn's test:<br>Left:<br>p>0.99 for WT vs. *Chd2*$^{WT/m}$<br>p=0.95 for WT vs. *Chd2*$^{m/m}$<br>Right:<br>p>0.99 for WT vs. *Chd2*$^{WT/m}$<br>p=0.0113 for WT vs. *Chd2*$^{m/m}$ |
| | D | 2-way ANOVA | WT: n=12 (F=7, M=5)<br>*Chd2*$^{WT/m}$: n=13 (F=6, M=7)<br>*Chd2*$^{m/m}$: n=12 (F=4, M=8) | p=0.0054 for the main effect of genotype | Dunnett's test:<br>p=0.0055 for WT vs. *Chd2*$^{WT/m}$<br>p=0.0146 for WT vs. *Chd2*$^{m/m}$<br><br>Statistical analysis of each trial is depicted in the graph |
| | F | Unpaired t-test | WT: n=9 from 3 mice<br>*Chd2*$^{m/m}$: n=9 from 3 mice | p=0.26 | |
| Fig. 3 | B | Kruskal-Wallis | WT: n=20 (F=11, M=9)<br>*Chd2*$^{WT/m}$: n=23 (F=11, M=12)<br>*Chd2*$^{m/m}$: n=15 (F=6, M=9) | Left:<br>p<0.0001<br>Right:<br>p<0.0001 | Dunn's test:<br>Left:<br>P=0.76 for WT vs. *Chd2*$^{WT/m}$<br>p<0.0001 for WT vs. *Chd2*$^{m/m}$<br>Right:<br>p=0.69 for WT vs. *Chd2*$^{WT/m}$<br>p<0.0001 for WT vs. *Chd2*$^{m/m}$ |
| | C | Kruskal-Wallis | WT: n=25 (F=11, M=14) | Left:<br>p<0.0001 | Dunn's test:<br>Left: |

| | | | | | |
|---|---|---|---|---|---|
| | | | *Chd2*$^{WT/m}$: n=27 (F=17, M=10)<br>*Chd2*$^{m/m}$: n=13 (F=6, M=7) | Right:<br>p<0.0001 | P=0.14 for WT vs. *Chd2*$^{WT/m}$<br>p<0.0001 for WT vs. *Chd2*$^{m/m}$<br>Right:<br>p=0.41 for WT vs. *Chd2*$^{WT/m}$<br>p<0.0001 for WT vs. *Chd2*$^{m/m}$ |
| | E | Kruskal-Wallis | Left:<br>WT: n=18 (F=11, M=7)<br>*Chd2*$^{WT/m}$: n=23 (F=13, M=10)<br>*Chd2*$^{m/m}$: n=17 (F=12, M=5)<br>Right:<br>WT: n=24 (F=14, M=10)<br>*Chd2*$^{WT/m}$: n=41 (F=21, M=20)<br>*Chd2*$^{m/m}$: n=21 (F=15, M=6) | Left:<br>p=0.008<br>Right:<br>p=0.0001 | Dunn's test:<br>Left:<br>p=0.185 for WT vs. *Chd2*$^{WT/m}$<br>p>0.99 for WT vs. *Chd2*$^{m/m}$<br>Right:<br>p<0.0001 for WT vs. *Chd2*$^{WT/m}$<br>p=0.14 for WT vs. *Chd2*$^{m/m}$ |
| | F | Two-way ANOVA | WT: n=10 mice in 5 pairs (F=2, M=8)<br>*Chd2*$^{WT/m}$: n=24 mice in 12 pairs (F=12, M=12) | Left:<br>p=0.27 for the main effect of genotype<br>p=0.028 for the interaction between time and genotypes<br>Right:<br>p=0.5 for the main effect of genotype<br>p=0.092 for the interaction between time and genotypes | Holm-Sidak's test:<br>Left:<br>p=0.99 for 0-5 min<br>p=0.99 for 5-10 min<br>p=0.019 for 10-15 min<br>Right:<br>p=0.67 for 0-5 min<br>p=0.9 for 5-10 min<br>p=0.038 for 10-15 min |
| Fig. 4 | B | Kcnj11: One-Way ANOVA | Kcnj11 (129X1/SvJ):<br>WT: n=3<br>*Chd2*$^{WT/m}$: n=3<br>*Chd2*$^{m/m}$: n=3<br>Kcnj11 (C57BL/6J):<br>WT: n=5<br>*Chd2*$^{WT/m}$: n=5<br>*Chd2*$^{m/m}$: n=3 | | Kcnj11 (129X1/SvJ): Holm-Sidak's test:<br>p=0.009 for WT vs. *Chd2*$^{WT/m}$<br>p=0.005 for WT vs. *Chd2*$^{m/m}$<br>Kcnj11 (C57BL/6J): Holm-Sidak's test:<br>p=0.036 for WT vs. *Chd2*$^{WT/m}$<br>p=0.036 for WT vs. *Chd2*$^{m/m}$ |
| | C | Kcnj11: One-Way ANOVA<br>Abcc8: Kruskal-Wallis | WT: n=3<br>*Chd2*$^{WT/m}$: n=3<br>*Chd2*$^{m/m}$: n=3 | | Kcnj11: Holm-Sidak's test:<br>p=0.16 for WT vs. *Chd2*$^{WT/m}$<br>p<0.0001 for WT vs. *Chd2*$^{m/m}$<br>Abcc8: Dunn's test:<br>p=0.59 for WT vs. *Chd2*$^{WT/m}$<br>p=0.022 for WT vs. *Chd2*$^{m/m}$ |
| | D | Kcnj11: One-Way ANOVA<br>Abcc8: Kruskal-Wallis | WT: n=3<br>*Chd2*$^{WT/m}$: n=4<br>*Chd2*$^{m/m}$: n=3 | | Kcnj11: Holm-Sidak's test:<br>p=0.47 for WT vs. *Chd2*$^{WT/m}$<br>p=0.0003 for WT vs. *Chd2*$^{m/m}$<br>Abcc8: Dunn's test:<br>p=0.41 for WT vs. *Chd2*$^{WT/m}$<br>p=0.014 for WT vs. *Chd2*$^{m/m}$ |
| Fig. 5 | B | One-Way ANOVA | WT: n=11 (F=5, M=6)<br>*Chd2*$^{WT/m}$: n=13 (F=3, M=10)<br>*Chd2*$^{m/m}$: n=10 (F=5, M=5) | p=0.0133 | Dunnett's test:<br>p=0.0096 for WT vs. *Chd2*$^{WT/m}$<br>p=0.0482 for WT vs. *Chd2*$^{m/m}$ |
| | C | Two-way ANOVA | WT: n=11 (F=5, M=6)<br>*Chd2*$^{WT/m}$: n=13 (F=3, M=10)<br>*Chd2*$^{m/m}$: n=10 (F=5, M=5) | p=0.0134 for the main effect of genotype | Holm-Sidak's test:<br>Delta:<br>p=0.0561 for WT vs. *Chd2*$^{WT/m}$<br>p=0.12 for WT vs. *Chd2*$^{m/m}$<br>Theta:<br>p=0.0265 for WT vs. *Chd2*$^{WT/m}$<br>p=0.0265 for WT vs. *Chd2*$^{m/m}$<br>Alpha: |

| | | | | |
|---|---|---|---|---|
| | | | | p=0.0238 for WT vs. *Chd2*$^{WT/m}$<br>p=0.0298 for WT vs. *Chd2*$^{m/m}$<br>Beta:<br>p=0.0187 for WT vs. *Chd2*$^{WT/m}$<br>p=0.05 for WT vs. *Chd2*$^{m/m}$<br>Gamma:<br>p=0.0497 for WT vs. *Chd2*$^{WT/m}$<br>p=0.5 for WT vs. *Chd2*$^{m/m}$ |
| E | Two-way ANOVA | WT: n=11 (F=5, M=6)<br>*Chd2*$^{WT/m}$: n=13 (F=3, M=10)<br>*Chd2*$^{m/m}$: n=10 (F=5, M=5) | p=0.42 for the main effect of genotype | Holm-Sidak's test:<br>Delta:<br>p=0.34 for WT vs. *Chd2*$^{WT/m}$<br>p=0.34 for WT vs. *Chd2*$^{m/m}$<br>Theta:<br>p=0.85 for WT vs. *Chd2*$^{WT/m}$<br>p=0.035 for WT vs. *Chd2*$^{m/m}$<br>Alpha:<br>p=0.17 for WT vs. *Chd2*$^{WT/m}$<br>p=0.17 for WT vs. *Chd2*$^{m/m}$<br>Beta:<br>p=0.44 for WT vs. *Chd2*$^{WT/m}$<br>p=0.54 for WT vs. *Chd2*$^{m/m}$<br>Gamma:<br>p=0.19 for WT vs. *Chd2*$^{WT/m}$<br>p=0.0366 for WT vs. *Chd2*$^{m/m}$ |
| F | Two-way ANOVA | WT: n=11 (F=5, M=6)<br>*Chd2*$^{WT/m}$: n=13 (F=3, M=10)<br>*Chd2*$^{m/m}$: n=10 (F=5, M=5) | p=0.15 for the main effect of genotype | Holm-Sidak's test:<br>Delta:<br>p=0.86 for WT vs. *Chd2*$^{WT/m}$<br>p=0.86 for WT vs. *Chd2*$^{m/m}$<br>Theta:<br>p=0.13 for WT vs. *Chd2*$^{WT/m}$<br>p=0.46 for WT vs. *Chd2*$^{m/m}$<br>Alpha:<br>p=0.0139 for WT vs. *Chd2*$^{WT/m}$<br>p=0.27 for WT vs. *Chd2*$^{m/m}$<br>Beta:<br>p=0.0486 for WT vs. *Chd2*$^{WT/m}$<br>p=0.66 for WT vs. *Chd2*$^{m/m}$<br>Gamma:<br>p=0.42 for WT vs. *Chd2*$^{WT/m}$<br>p=0.2 for WT vs. *Chd2*$^{m/m}$ |
| H | Log-rank | WT: n=6 (F=5, M=1)<br>*Chd2*$^{WT/m}$: n=6 (F=1, M=5)<br>*Chd2*$^{m/m}$: n=3 (F=1, M=2) | p=0.0497 for WT vs. *Chd2*$^{WT/m}$<br>p=0.097 for WT vs. *Chd2*$^{m/m}$ | |
| I | Chi-square | WT: n=6 (F=5, M=1)<br>*Chd2*$^{WT/m}$: n=6 (F=1, M=5)<br>*Chd2*$^{m/m}$: n=3 (F=1, M=2) | p=0.0209 for WT vs. *Chd2*$^{WT/m}$<br>p=0.0177 for WT vs. *Chd2*$^{m/m}$ | |
